# Characterisation of native human pancreatic mesenchymal stromal cells in type 1 diabetes

**DOI:** 10.1101/2025.03.05.641596

**Authors:** Rebecca E. Dewhurst-Trigg, Jocelyn Atkins, Noel G. Morgan, Martin Eichmann, Sarah J. Richardson, Chloe L. Rackham

## Abstract

**Aims/hypothesis:** Culture-expanded mesenchymal stromal cells (MSCs) reduce immune cell activation and improve islet functional survival. However, little is known about native human pancreatic MSCs (npMSCs) in health or how they are altered in type 1 diabetes. Here, we determined the number, density and islet-protective phenotype of npMSCs in situ in control individuals and those with type 1 diabetes.

**Methods:** Multiplex immunohistochemistry was used to identify npMSCs (CD90^+^/CD105^+^/CD73^+^/CD31^-^/CD45^-^/CD34^-^) in human pancreas sections from 38 donors (Network for Pancreatic Organ Donors with Diabetes and Exeter Archival Diabetes Biobank). Donors were categorised as either < 13 years at type 1 diabetes diagnosis (*n* = 8) or ≥ 13 years at type 1 diabetes diagnosis (*n* = 11) or were age- and sex-matched individuals without diabetes. Consecutive sections were immunostained with antisera against insulin, glucagon, and the established islet-protective and immunomodulatory factors Annexin A1 (ANXA1) and indoleamine 2,3-Dioxygenase-1. Whole-slide scans were acquired and npMSCs either inside or at the periphery (within 10 µm) of islets were quantified on an individual-islet basis. We identified 53,375 npMSCs and performed an analysis of 26,376 individual islets. Culture-expanded MSCs were exposed to cytokines and viability and proliferation was assessed by flow cytometry.

**Results:** npMSC were identified in situ in the human pancreas where they wrap around the islet periphery in an expected spindle-like morphology. ANXA1 was expressed by 33.2% of npMSCs and was expressed constitutively among individuals with or without diabetes. The density of both intraislet npMSCs and npMSCs within 10 µm of the islet periphery was increased for insulin-containing islets in individuals with type 1 diabetes compared to individuals without diabetes (*p* < 0.001). npMSC density within 10 µm of the islet periphery was preferentially increased in individuals ≥ 13 years at type 1 diabetes diagnosis compared to individuals < 13 years at type 1 diabetes diagnosis (*p* < 0.001). npMSC density was reduced around insulin-deficient islets compared to insulin-containing islets in individuals with diabetes (*p* < 0.001), consistent with an islet-protective role for npMSCs. Exposure of culture-expanded MSCs to an aggressive cytokine combination led to increased cell death and reduced proliferation.

**Conclusion/interpretation:** npMSCs express ANXA1 constitutively suggesting an islet-protective role in health. The density of npMSCs was increased around insulin-containing islets and lost around insulin-deficient islets in individuals with type 1 diabetes which aligns with this hypothesis. npMSC density at the periphery of insulin-containing islets was preferentially higher in individuals with later onset type 1 diabetes, correlating with a less intense immune cell infiltration. The reduced ability of npMSCs to survive in the more intense pro-inflammatory environment around islets in younger onset type 1 diabetes may contribute to the rapid rate of beta cell loss in these individuals.

**Research in context:** 

**What is already known about the subject?:** • Culture-expanded mesenchymal stromal cells (MSCs) reduce immune cell activation and improve islet functional survival.

• Therapeutically administered MSCs migrate to sites of injury and preserve endogenous C-peptide in individuals with newly diagnosed type 1 diabetes.

**What is the key question?:** • Is the number, distribution and/or cytoprotective and immunomodulatory phenotype of native pancreatic MSCs (npMSCs) altered in type 1 diabetes?

**What are the new findings?:** • npMSCs express typical markers of isolated, culture-expanded MSCs and their number and density is increased within and at the periphery of insulin-containing islets in individuals with type 1 diabetes.

• npMSCs are more abundant at the periphery of islets in individuals with later onset type 1 diabetes, correlating with a less intense immune cell infiltration.

• MSCs are less able to survive an aggressive inflammatory environment such as that arising from the enhanced islet immune cell infiltration in individuals with younger onset type 1 diabetes

**How might this impact on clinical practice in the foreseeable future?:** • Therapeutically administered MSCs are likely to be more effective at preserving endogenous beta cell mass in individuals with later onset type 1 diabetes where MSCs have a greater ability to survive and exert their therapeutic functions.

## Introduction

Mesenchymal Stromal Cells (MSCs) form the focus of attention in an increasing number of translational research fields, including diabetes [1–4], because of their strong immunomodulatory [5, 6] and regenerative [7–10] properties. We have begun to reveal the therapeutic mechanisms by which isolated and “culture-expanded” MSCs (exogenous MSCs), derived from clinically relevant tissue sources (including bone marrow, adipose and pancreas) modulate islet function and survival [11–15], as well as immune cell activity [5, 16–18]. Intravenous infusion of culture-expanded MSCs preserves endogenous C-peptide in individuals with newly diagnosed type 1 diabetes [1, 2]. MSCs respond to specific cues within their microenvironment and migrate to sites of injury, including pancreatic islets [19], to exert their therapeutic actions.

Several studies have demonstrated that (culture-expanded) pancreas-derived MSCs have similar immunomodulatory and therapeutic potential to archetypal bone marrow MSCs [7, 8, 20–23]. These conclusions have been drawn based on studies using MSCs isolated initially from the pancreas, then expanded in culture and characterised according to their differentiation potential and surface expression of a panel of defined markers. Specifically, human culture-expanded MSCs, including those derived from pancreas [20, 24–26], are immunopositive for several surface antigens (CD73, CD90/Thy-1, CD105/Endoglin, whilst they lack hematopoietic (CD34, CD45) and endothelial cell markers (CD31) [27, 28]. To our knowledge, a detailed immunohistological characterisation of native human pancreatic MSCs (npMSCs) in situ has not been completed in individuals either with or without type 1 diabetes. Accordingly, little is known about the physiological roles of npMSCs nor how this may be altered in type 1 diabetes.

No single phenotypic marker can distinguish MSCs from other stromal or perivascular cell types. Multiplex immunohistochemistry methods therefore offer a robust approach for high-resolution, simultaneous detection of multiple antigens within a single tissue section and provides a suitable methodology for the detection and characterisation of npMSCs in situ. We have utilised multiplex immunohistochemistry to identify npMSCs (CD73^+^, CD90^+^, CD105^+^, CD31^-^, CD34^-^, CD45^-^) in human pancreas sections from 38 donors (Network for Pancreatic Organ Donors with Diabetes (nPOD) and Exeter Archival Diabetes Biobank (EADB)) with short-duration type 1 diabetes (≤ 2 years) or similarly aged donors without type 1 diabetes. Multiplex immunohistochemistry is particularly advantageous in studies using rare and finite tissue resources, such as pancreas tissue available for research in young individuals with short-duration type 1 diabetes. To maximise insights into type 1 diabetes disease mechanisms and progression, the conservation of these valuable and irreplaceable tissue samples is essential across the research community. Accordingly, this study employed the Opal-tyramide signal amplification immunostaining method, whole-slide scanning (PhenoImager HT) and the quantitative digital pathology platform HALO, to maximise data generation from each tissue section. This approach enabled the simultaneous detection of six distinct tissue antigens per section and data analysis at the individual islet level. Understanding how the number, distribution and islet-protective phenotype of npMSCs is altered in type 1 diabetes is of critical importance as these cells become a therapeutic option for intervention in type 1 diabetes pathogenesis and progression.

## Methods

### Human tissue

Formalin- or mercuric chloride-fixed, paraffin-embedded pancreas tissue sessions were obtained from within the EADB and nPOD collections. Ethical permission from the West of Scotland Research Ethics Committee (ref: 20/WS/007; IRAS Project ID: 283620) and from the nPOD tissue prioritisation committee, University of Florida (Project ID: 24-013) was obtained for the use of EADB and nPOD samples, respectively.

Analyses were performed using donors with short-duration (≤ 2 years) type 1 diabetes who were < 13 years at type 1 diabetes diagnosis or ≥ 13 years at type 1 diabetes diagnosis. Control individuals were sex-matched donors without type 1 diabetes of similar age and BMI. The cut-off for defining two groups according to age of diagnosis has been informed by previous studies in which individuals with type 1 diabetes were categorised as type 1 diabetes endotype 1 (T1DE1) or type 1 diabetes endotype 2 (T1DE2), according to distinct histological phenotypes (CD20Hi/Lo) and extent of proinsulin processing [29]. Individuals diagnosed < 13 years often present with a more aggressive disease associated with increased immune cell infiltration and greater loss of residual beta cells in the pancreas [29, 30]. We acknowledge that the current report does not directly categorise type 1 diabetes individuals according to these distinct histological phenotypes to avoid unnecessary tissue utilisation. Instead, it used an age at diagnosis cut-off of 13 years to align with the T1DE2 categorisation on the understanding that most younger subjects will then be defined as T1DE1. Thus, for the current report, our type 1 diabetes subcategorisation is not absolutely endotype-specific but, as a surrogate, it stratifies specific groups of individuals by virtue of their age at diagnosis. Individuals were included in the current study from the EADB (*n* = 3, < 13 years at type 1 diabetes diagnosis; *n* = 3, < 13 years without diabetes; *n* = 3, ≥ 13 years at type 1 diabetes diagnosis; *n* = 3, ≥ 13 years without diabetes) and nPOD (*n* = 5, < 13 years at type 1 diabetes diagnosis; *n* = 5, < 13 years without diabetes; *n* = 8, ≥ 13 years at type 1 diabetes diagnosis; *n* = 8, ≥ 13 years without diabetes) collections. Individual donor characteristics are detailed in Electronic Supplementary Material (ESM) Table 1. Age was similar between individuals < 13 years with type 1 diabetes and without diabetes (mean ± SEM; 6.80 ± 1.54 years vs 6.13 ± 1.01 years, respectively; t(14) = 0.3656, *p* = 0.72) and between individuals ≥ 13 years with type 1 diabetes and without diabetes (19.35 ± 1.44 years vs 17.75 ± 1.34 years, respectively; t(20) = 0.8085, *p* = 0.43). Donors of European descent were selected to ensure that any alterations in npMSCs were not due to the emerging evidence of heterogeneity of type 1 diabetes between ethnic groups (reviewed in [31]). For nPOD donors where information was available, pancreas sections were taken from the pancreas tail and block numbers were matched for individuals with and without diabetes.

**Table 1.**
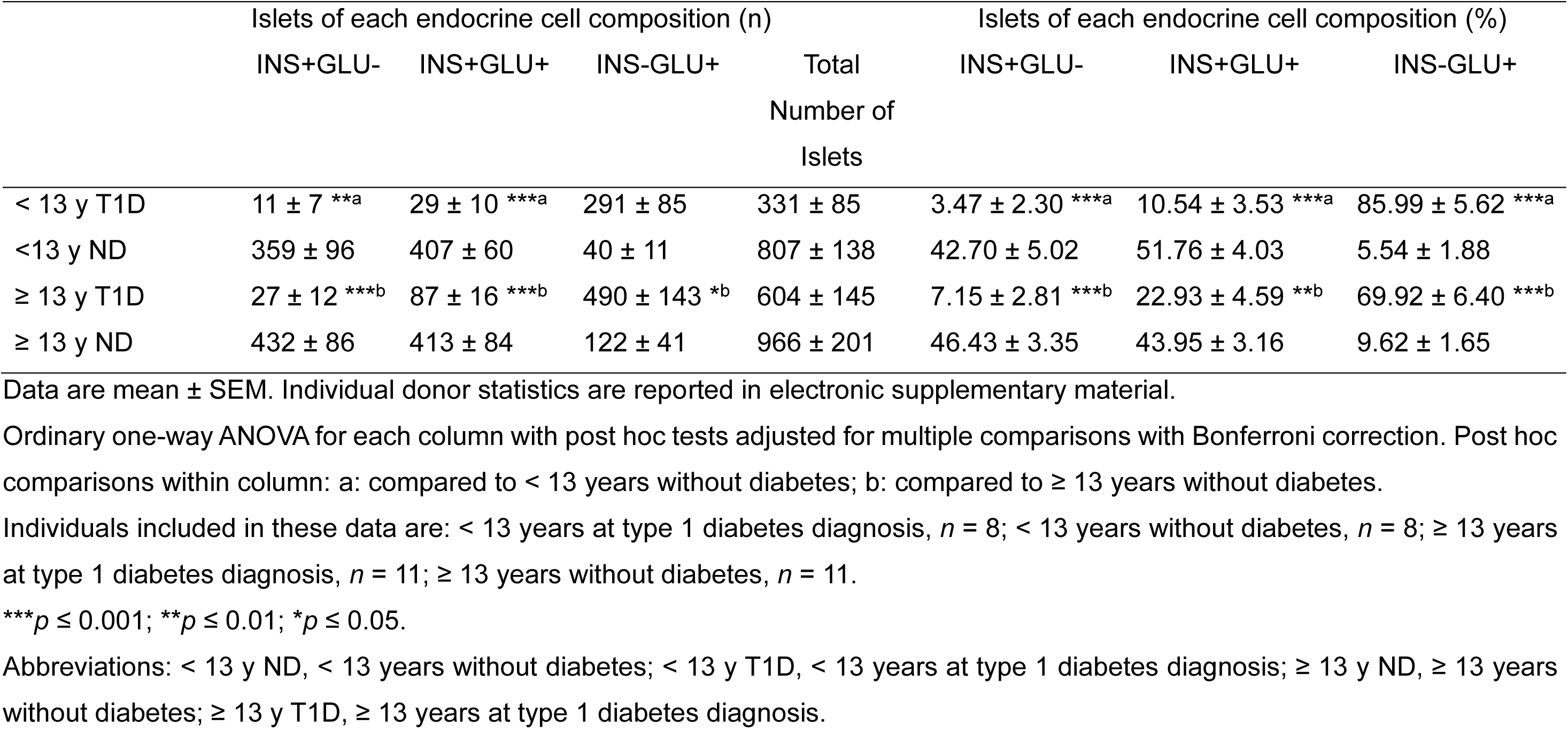
Average number and percentage of islets comprising different endocrine cell compositions among individuals with and without type 1 diabetes from the EADB and nPOD collections.

### Multiplex Immunohistochemistry

Four-micron human pancreas, tonsil, intestine and placenta sections from the EADB collection were used for initial antibody and subsequent Opal 6-plex panel optimisation. The Opal multiplex panel optimisation is summarised in ESM Fig. 1. The final 6-plex panels to identify npMSCs and islet hormone and MSC islet-protective factors are detailed in ESM Tables 2 and 3, respectively.

**Fig. 1.**
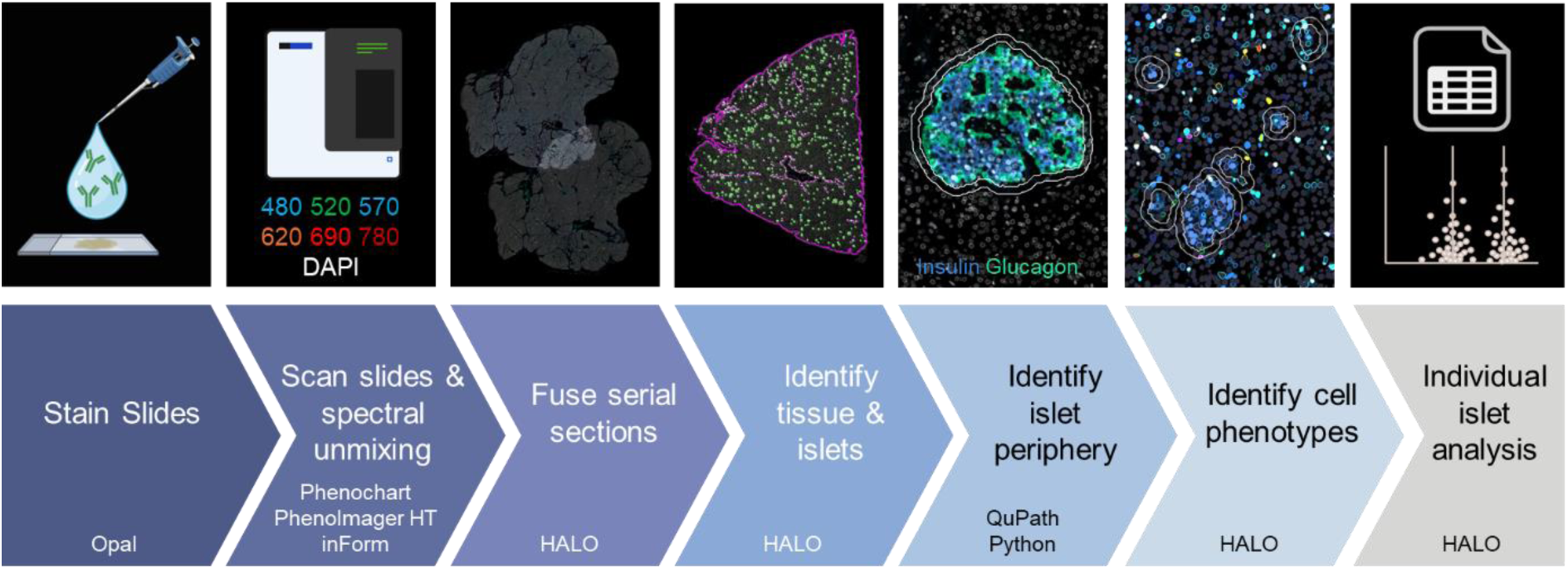
Optimised workflow for the generation of quantitative data, on an individual-islet basis, for the number and density of npMSCs associated with islets of different endocrine cell compositions and size. Serial sections were immunostained with our optimised 6-plex Opal panel to identify npMSCs or islet hormones and MSC-derived islet-protective factors. Slides were scanned at x40 magnification using the MOTiF workflow on a PhenoImager HT and multiplex images were spectrally unmixed. Unmixed whole-slide scans were loaded into HALO and serial sections registered and fused. A Random Forest classifier was used to identify glass, tissue, and islets that were more than 170 µm^2^ (magenta annotation shows pancreas outline, and green annotations show islets). Islet annotations were imported into QuPath and expanded 10 µm or until they reached another expanding annotation (white annotations show the islet periphery, and the islet annotation expanded by 10 µm). Positive staining thresholds were set for all antigens and cell phenotypes identified. Data analysis was performed on each individual islet and expanded islet annotation (islet periphery data determined by expanded islet minus islet data). Donor ID: nPOD 6271, EADB 12426, EADB E386, nPOD 6488. Figure created using BioRender.

Following tissue rehydration, multiplex immunostaining was performed using the Opal 6-plex manual detection kit (Akoya Biosciences). Heat induced epitope retrieval (HIER) was performed using a pressure cooker. HIER buffers were made in-house containing either 0.01M citrate (Merck), pH 6, or 10 mM TrisBase (Merck) and 1 mM EDTA (Merck), pH 9 and were specific to each antigen (see ESM Tables 2 and 3 for details).

One section per donor was immunostained with antibodies against CD90, CD105, CD73, CD31, CD34 and CD45 to identify npMSCs. A serial section was then immunostained with antibodies against CD90, Indoleamine 2,3-dioxygenase 1 (IDO1), Annexin A1 (ANXA1), CD45, insulin and glucagon to identify islets and MSC-derived islet-protective factors. Antigens were visualised using the Opal fluorophores: 480, 520, 570, 620, 690 and 780, and sections were counterstained with 4’,6-diamidino-2-phenylindole (DAPI; Life Technologies). Once the immunostaining was complete, sections were mounted using ProLong Diamond Antifade mounting medium (Fisher Scientific).

### Image acquisition and processing

Slides were scanned at x40 magnification using the MOTiF workflow on a PhenoImager HT (Akoya Biosciences) within two weeks of mounting the tissue. Exposure times were determined as the mean time following auto-exposure at several positions across the tissue. Exposure times were set separately for each tissue biobank (EADB and nPOD) and remained consistent across all sections within each biobank. Spectral unmixing was performed in InForm using the Akoya synthetic library. Autofluorescence subtraction was adjusted individually for sections from EADB due to variations in innate tissue autofluorescence across sections. Autofluorescence subtraction remained consistent for all nPOD sections where autofluorescence remained similar across sections.

### Image quantification

Image quantification was performed in HALO (v3.6, Indica Labs). Serial sections from each donor were registered and fused. Then, a random forest classifier was used to identify pancreas tissue and islets (based on positive insulin and/or glucagon staining) that were larger than the estimated area of at least one cell (170 µm^2^). Islet annotations were generated from the classifier and exported from HALO as a .GeoJSON file. All islet annotations were simultaneously expanded outwards in QuPath (v0.5.1) by 10 µm or until they reached another expanding annotation to identify the islet periphery. This avoided any overlapping annotation areas, which ensured that cells were not counted twice during individual islet analysis. Each islet annotation was assigned a unique identifier in Python and then re-imported into HALO for analysis. Quantitative analysis of cell populations was performed using HighPlex FL (v4.2.14, HALO, Indica Labs). First, cells were identified using DAPI and positive cytoplasmic and nuclear staining thresholds were set for all antigens. Staining thresholds were adjusted for each donor individually. Next, cell phenotypes were determined and npMSCs were identified as CD90^+^, CD73^+^, CD105^+^, CD31^-^, CD34^-^, CD45^-^ cells. HighPlex FL analysis was run in batch across all annotation layers to give individual islet data, including intraislet data and islet periphery (10 µm outside of the islet) data (calculated by expanded islet minus islet annotation).

Prior to analysis, individual islets were categorised into one of three categories based on their endocrine cell composition: insulin-positive glucagon-negative (INS+GLU-), insulin-positive glucagon-positive (INS+GLU+), or insulin-negative glucagon-positive (INS-GLU+). Islets were categorised as positive for insulin or glucagon if they contained one or more positive cell for the respective marker. Islets were further categorised by size by calculating the base-2 logarithm of the ratio between the measured islet area and the estimated individual cell area (170 µm^2^), then rounded down to the nearest integer. Representative images of islets from each size category are shown in ESM Fig. 2.

**Fig. 2.**
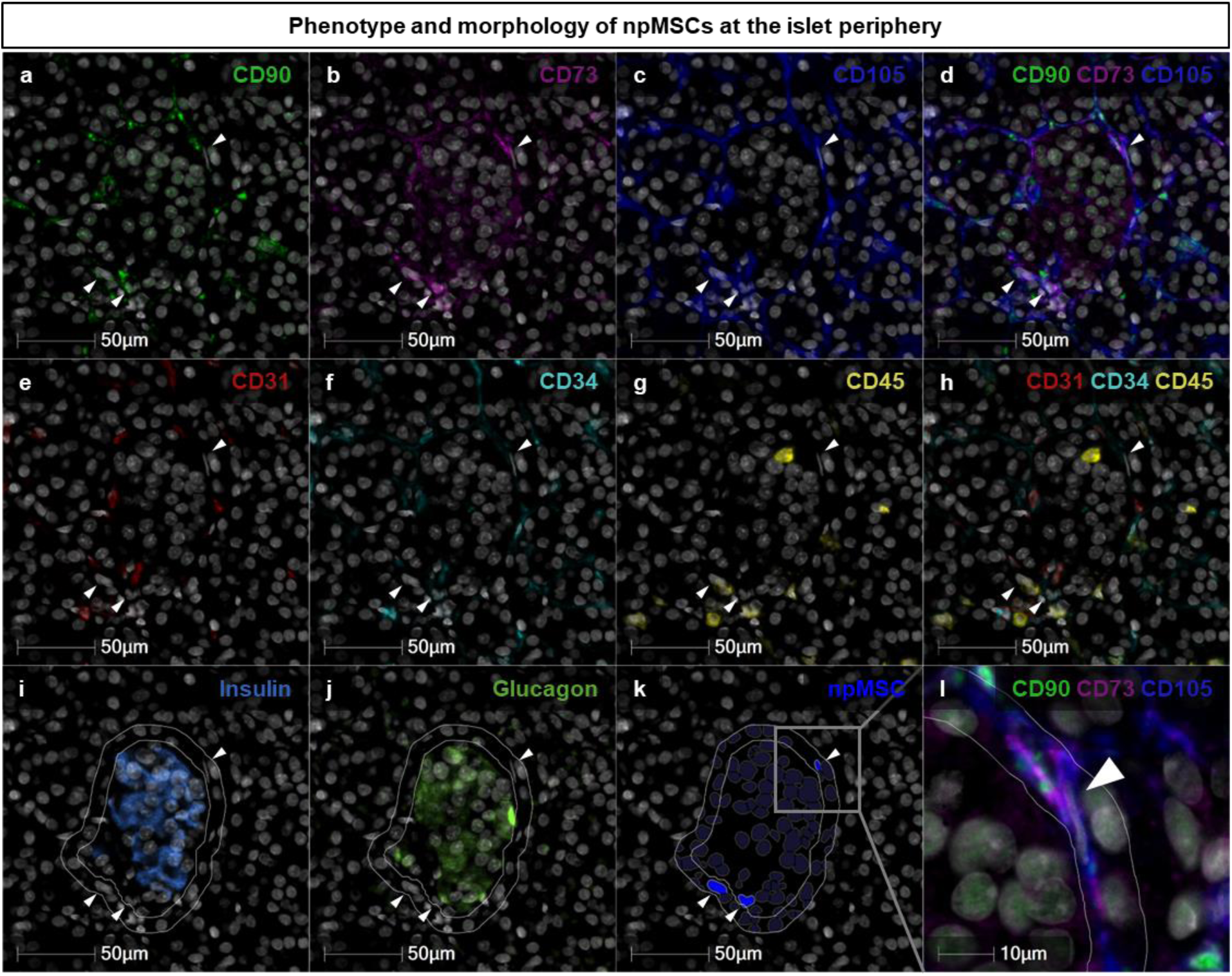
Phenotype and morphology of npMSCs at the islet periphery. npMSCs (shown by white arrows, Fig. 2a-l) at the islet periphery were identified in HALO as CD90+ (a), CD73+ (b), CD105+ ((c); overlay of positive markers, (d)), CD31- (e), CD34- (f), CD45- ((g); overlay of negative markers, (h)). Islets were identified by insulin (i) and glucagon (j) immunostaining (inner white annotation) and the islet annotation expanded 10 µm or until another annotation was reached (outer white annotation). Next, npMSCs were quantified using the HALO HighPlex FL package (k). A magnified micrograph of k is shown in l (grey box shows magnified area) demonstrating that npMSCs appear to be wrapped around the islet and display an elongated spindle-shaped morphology (l). Micrographs used in the representative figure were adjusted to optimise contrast and visibility without altering the underlying image data or quantification. Adjustments were made to the ‘Black In’, ‘White In’ and ‘Gamma’ settings. Donor ID: Individual ≥ 13 years at type 1 diabetes diagnosis, 6563, nPOD.

Our optimised workflow for the generation of quantitative data on an individual-islet basis for the number, density and islet-protective phenotype of npMSCs associated with islets of different endocrine cell compositions and size is shown in Fig. 1.

### MSC viability and proliferation following cytokine exposure in vitro

Human adipose tissue-derived MSCs (StemPro Human Adipose-derived Stem Cells; female donor 51 years, Fisher Scientific; verified mycoplasma-free in-house, MycoAlert Plus, Lonza Biosciences) were labelled with 0.5 mM CellTrace Far Red (Invitrogen) according to manufacturer’s recommendations to enable quantification of proliferation and seeded into 6 well plates at 75,000 cells/well. MSCs were cultured in DMEM supplemented with 10% (vol/vol) FBS, 100 U/mL penicillin, 100 µg/mL streptomycin and 2 mM L-glutamine. MSCs were cultured at 37°C in a humidified environment containing 5% CO_2_. MSCs were left to adhere overnight and then cultured for three or seven days in either normal medium (no cytokine control) or normal medium containing either: IFN-α alone; IFN-γ + IL-1β; or IFN-γ + IL-1β + TNF-α. The concentration of each cytokine was: IFN-α (1000 U/mL, PBL Assay Science), IFN-γ (1000 U/mL, PeproTech), IL-1β (50 U/mL, PeproTech) and TNF-α (1000 U/mL, PeproTech). Media was refreshed on day three of cytokine exposure. To prepare samples for analysis, supernatant was collected, and cells were detached with Accutase. Cells were washed in cold PBS then 0.5% FBS in PBS and resuspended in 600 µL 0.5% FBS in PBS. To quantify MSC viability, one drop of Sytox Green (Fisher Scientific) was added 5 min before acquisition by Flow Cytometry (Attune, ThermoFisher Scientific). Flow cytometry analysis of MSC viability and proliferation was performed in FlowJo (v10.8.1) using the gating strategy shown in ESM Fig. 3. The percentage of divided cells was calculated using the FlowJo proliferation algorithm to determine proliferation of CellTrace-labelled cells.

**Fig. 3.**
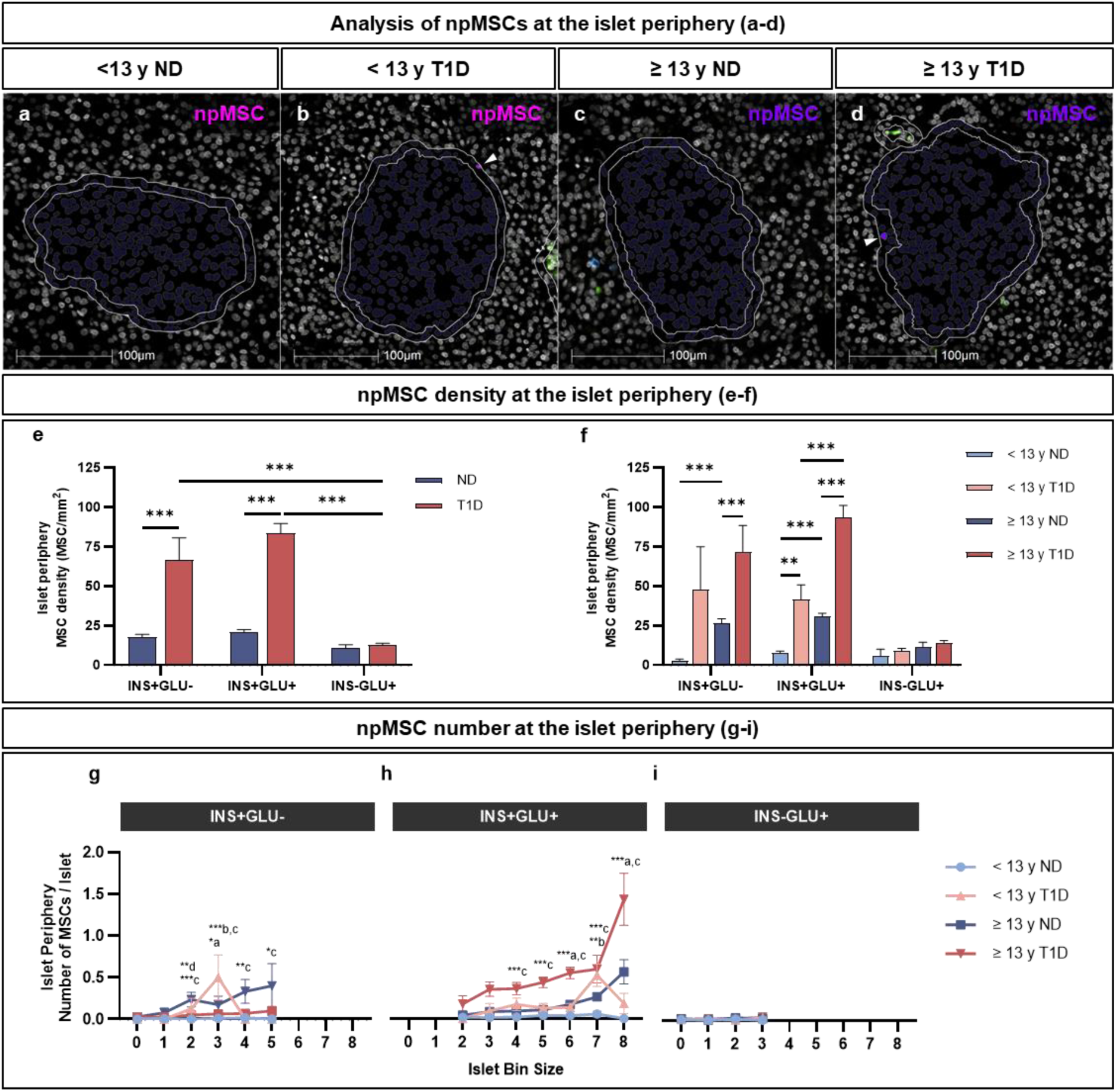
The density and number of npMSCs within a 10 µm region at the periphery of islets comprising different endocrine cell compositions and size. Representative quantification of npMSCs (HighPlex FL package, HALO) at the islet periphery are shown in Fig. 3a-d (npMSCs highlighted by white arrows). For Fig. 3e-f, data derived from *n* = 26,376 individual islets from 38 individuals (*n* = 8 individuals < 13 years at type 1 diabetes diagnosis and *n* = 11 individuals ≥ 13 years at type 1 diabetes diagnosis (*n* = 19 individuals with type 1 diabetes); *n* = 8 individuals < 13 years without diabetes and *n* = 11 individuals > 13 years without diabetes (*n* = 19 individuals without diabetes)). For Fig. 3o-p, islet bin sizes containing < 5 islets, or bin sizes devoid of data for one or more group were excluded; data derived from *n* = 23,290 individual islets. Bars represent mean ± SEM. Two-way ordinary ANOVA with post hoc tests adjusted for multiple comparisons with Bonferroni correction. Post hoc tests in Fig. 3e: islets of the same endocrine cell composition; diabetes or no diabetes across islets of different endocrine cell compositions. Post hoc tests in Fig. 3f: type 1 diabetes group and age-matched individuals without diabetes; type 1 diabetes and age of diagnosis; individuals without diabetes. Comparisons in Fig. 3e-f shown with a horizontal black line between the two compared groups. Post hoc comparisons within each islet bin size in Fig. 3g-h: < 13 years at type 1 diabetes diagnosis vs ≥ 13 years at type 1 diabetes diagnosis (in-figure label ‘a’); < 13 years at type 1 diabetes diagnosis vs < 13 years without diabetes (in-figure label ‘b’); ≥ 13 years at type 1 diabetes diagnosis vs ≥ 13 years without diabetes (in-figure label ‘c’); < 13 years without diabetes vs ≥ 13 years without diabetes (in-figure label ‘d)’. \*\*\**p* ≤ 0.001; \*\**p* ≤ 0.01; \**p* ≤ 0.05. Donor ID: nPOD 6407 (a), nPOD 6578 (b), nPOD 6333 (c), nPOD 6362 (d). Abbreviations: < 13 y ND, < 13 years without diabetes; < 13 y T1D, < 13 years at type 1 diabetes diagnosis; ≥ 13 y ND, ≥ 13 years without diabetes; ≥ 13 y T1D, ≥ 13 years at type 1 diabetes diagnosis.

### Statistical analysis

Quantitative data on an individual-islet basis were generated from whole-slide scans using HALO. Raw data were exported to Excel, and descriptive statistics (including mean, SD, SEM and n) generated with RStudio (v4.4.0). Preliminary code generation was facilitated using an artificial intelligence language model (ChatGPT). All generated code was reviewed, validated and refined manually to ensure accuracy and suitability for this study. Statistical analysis and graphical representation of these data were performed in GraphPad Prism (v10) and included unpaired t test, one-way ANOVA and two-way ANOVA, as indicated in each figure legend. Where appropriate, post hoc analyses for ANOVAs were performed, and were adjusted for multiple comparisons with the Bonferroni correction. *p* values are reported as: \*\*\**p* ≤ 0.001; \*\**p* ≤ 0.01; \**p* ≤ 0.05. All data are reported as mean ± SEM.

## Results

### Islet endocrine cell composition among individuals with and without type 1 diabetes

A total of 26,376 individual islets were identified across the 38 individuals with or without type 1 diabetes included in this study. The number and proportion of islets comprising each endocrine cell composition (INS+GLU+, INS+GLU-, and INS-GLU+) is shown as an average per group in Table 1 and for individual donors in ESM Table 4. Our 2D data from individuals without diabetes align with recent observations made in adult pancreas using 3D imaging technologies, which highlight that a significant proportion (approximately 50 %) of insulin-containing islets are devoid of glucagon [32].

In line with the classification of individuals with type 1 diabetes into two groups based on age at diagnosis and therefore aggressiveness in beta cell destruction, the proportion of INS+GLU+ islets tended to be higher in individuals ≥ 13 years at diagnosis of type 1 diabetes compared to individuals < 13 years at diagnosis of type 1 diabetes (mean ± SEM; 22.93 ± 4.59 % vs 10.54 ± 3.53 %, respectively; unpaired t test: t(17) = 2.002, *p* = 0.06). This aligns with previous observations demonstrating a higher proportion of insulin-containing islets in individuals with T1DE2 compared to T1DE1 [29, 30]. The identification of islets according to the three defined endocrine cell compositions allowed us to investigate whether npMSCs were preferentially associated with islets of a particular endocrine cell composition.

### An extensive 6-plex panel is required to accurately identify npMSCs in situ

MSCs share phenotypic markers with endothelial cells, pericytes, fibroblasts and stellate cells. Thus, no single phenotypic marker can distinguish MSCs from other stromal or perivascular cell types. We evaluated whether a simplified 3-plex panel (CD90^+^, CD105^+^, CD31^-^) could accurately phenotype npMSCs in situ, or whether a more comprehensive 6-plex panel was required. CD90 and CD105 are widely used as positive MSC markers, however they are also expressed by endothelial cells. To address this, we included the endothelial cell marker CD31 as a negative MSC marker. The 6-plex panel included CD73 as an additional positive MSC marker, as well as CD34 (endothelial cell and hematopoietic stem cell marker) and CD45 (pan-immune cell marker) as additional negative MSC markers (full 6-plex panel: CD90^+^, CD73^+^ CD105^+^, CD31^-^, CD34^-^, CD45^-^).

Whole pancreas MSC density across EADB and nPOD donors was markedly overestimated when phenotyped using a core 3-plex (CD90^+^, CD105^+^, CD31^-^), compared to the comprehensive 6-plex panel (CD73^+^, CD90^+^, CD105^+^, CD31^-^, CD34^-^ , CD45^-^ (mean ± SEM; 123 ± 19 vs 14 ± 3 MSCs/mm^2^, respectively; t(74)=5.647, *p* < 0.001)). This validates the importance of characterising npMSCs using a 6-plex panel, to ensure cell types of distinct but similar phenotypes are excluded from the npMSC population. Using the comprehensive 6-plex panel, a total of 53,375 npMSCs (CD73^+^, CD90^+^, CD105^+^, CD31^-^, CD34^-^, CD45^-^) were identified across the 38 individuals included in this study. The npMSC population defined by the comprehensive 6-plex panel was further analysed to determine whether their number, density and/or islet-protective phenotype was altered in type 1 diabetes.

### Whole pancreas npMSC density is similar in individuals with and without type 1 diabetes

The density of npMSCs across the total pancreas area including acinar, islets, ducts and vessels was variable between individuals both with and without type 1 diabetes (range 0 – 63 and 1 – 94 npMSCs/mm^2^, respectively), as expected given the established migratory capacity of MSCs [19]. Together, the density of npMSCs across the whole pancreas area was similar for individuals with and without type 1 diabetes (14.70 ± 3.98 vs 14.07 ± 5.17 npMSCs/mm^2^, respectively; t(36) = 0.0966, *p* = 0.92). When separated by age, total pancreas npMSC density was similar between individuals with or without type 1 diabetes (one-way ANOVA; F (3, 34) = 0.9442, *p* = 0.43).

### Detection of npMSCs in close proximity to pancreatic islets

npMSCs, defined as immunopositive for CD90, CD73 and CD105 (Fig. 2a-d) and immunonegative for CD31, CD34 and CD45 (Fig. 2e-h), that were associated (either inside or within 10μm of the outside of the islet) with islets (Fig. 2i-j) were identified (Fig. 2a-k and ESM Fig. 4). Across all 38 donors, 3796 npMSCs (7 % of total pancreas MSCs) were associated with islets. A total of 1699 npMSCs at the islet periphery (within 10 µm of the outside of the islet), and 2097 intraislet npMSCs were identified. npMSCs appear to be wrapped around the islet periphery and display an elongated, spindle-shaped morphology, as expected (Fig. 2.l).

**Fig. 4.**
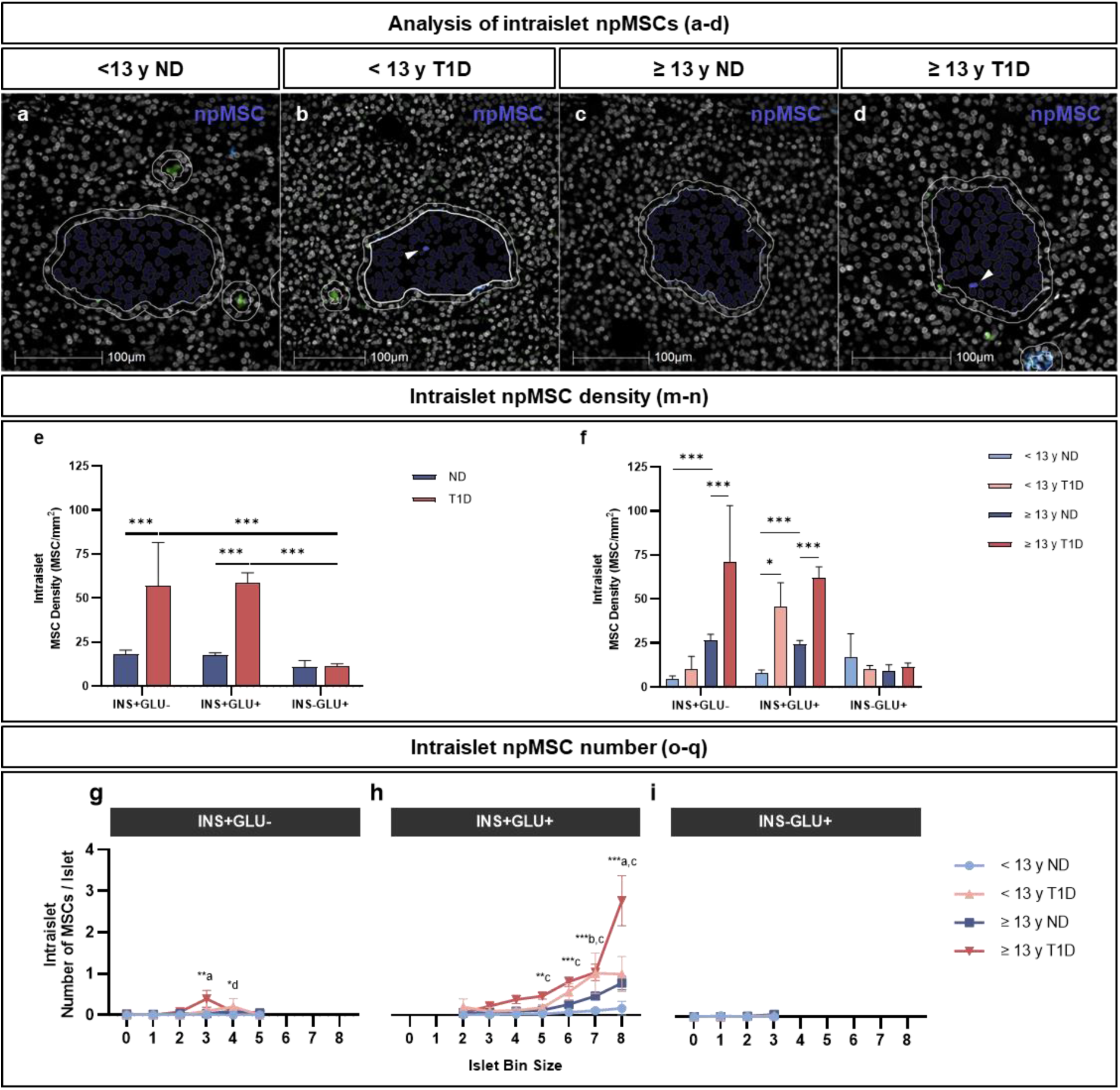
The density and number of intraislet npMSCs of islets comprising different endocrine cell compositions and size. Intraislet npMSCs (shown by white arrows, Fig. 4a-d) within INS+GLU+ islets are shown. Immunostaining for insulin (blue) and glucagon (green) outside of the analysed islets in a-d is shown. For Fig. 4e-f, data derived from *n* = 26,376 individual islets from 38 individuals (*n* = 8 individuals < 13 years at type 1 diabetes diagnosis and *n* = 11 individuals ≥ 13 years at type 1 diabetes diagnosis (*n* = 19 individuals with type 1 diabetes); *n* = 8 individuals < 13 years without diabetes and *n* = 11 individuals > 13 years without diabetes (*n* = 19 individuals without diabetes)). For Fig. 4g-I, islet bin sizes containing < 5 islets, or bin sizes devoid of data for one or more group were excluded; data derived from *n* = 23,290 individual islets. Bars represent mean ± SEM. Two-way ordinary ANOVA with post hoc tests adjusted for multiple comparisons with Bonferroni correction. Post hoc tests in Fig. 4e: islets of the same endocrine cell composition; diabetes or no diabetes across islets of different endocrine cell compositions. Post hoc tests in Fig. 4f: type 1 diabetes group and age-matched individuals without diabetes; type 1 diabetes and age of onset; individuals without diabetes. Comparisons in Fig. 4e-f shown with a horizontal black line between the two compared groups. Post hoc comparisons within each islet bin size in Fig. 4g-i: < 13 years at type 1 diabetes diagnosis vs ≥ 13 years at type 1 diabetes diagnosis (in-figure label ‘a’); < 13 years at type 1 diabetes diagnosis vs < 13 years without diabetes (in-figure label ‘b’); ≥ 13 years at type 1 diabetes diagnosis vs ≥ 13 years without diabetes (in-figure label ‘c’); < 13 years without diabetes vs ≥ 13 years without (in-figure label ‘d)’. \*\*\**p* ≤ 0.001; \*\**p* ≤ 0.01; \**p* ≤ 0.05. Donor ID: nPOD 6407 (a); nPOD 6578 (b), nPOD 6333 (c); nPOD 6362 (d). Abbreviations: < 13 y ND, < 13 years without diabetes; < 13 y T1D, < 13 years at type 1 diabetes diagnosis; ≥ 13 y ND, ≥ 13 years without diabetes; ≥ 13 y T1D, ≥ 13 years at type 1 diabetes diagnosis.

Islets that had an associated npMSC were defined as islets with one or more npMSC either within 10 µm of the islet periphery or within the islet itself (intraislet). The percentage of INS+GLU+ islets that had one or more associated npMSC was higher in individuals with type 1 diabetes (≥ 39.5 %) either < 13 years or ≥ 13 years at diagnosis, compared to respective age-matched individuals without diabetes (≤ 20.3%; ESM Fig. 5).

**Fig. 5.**
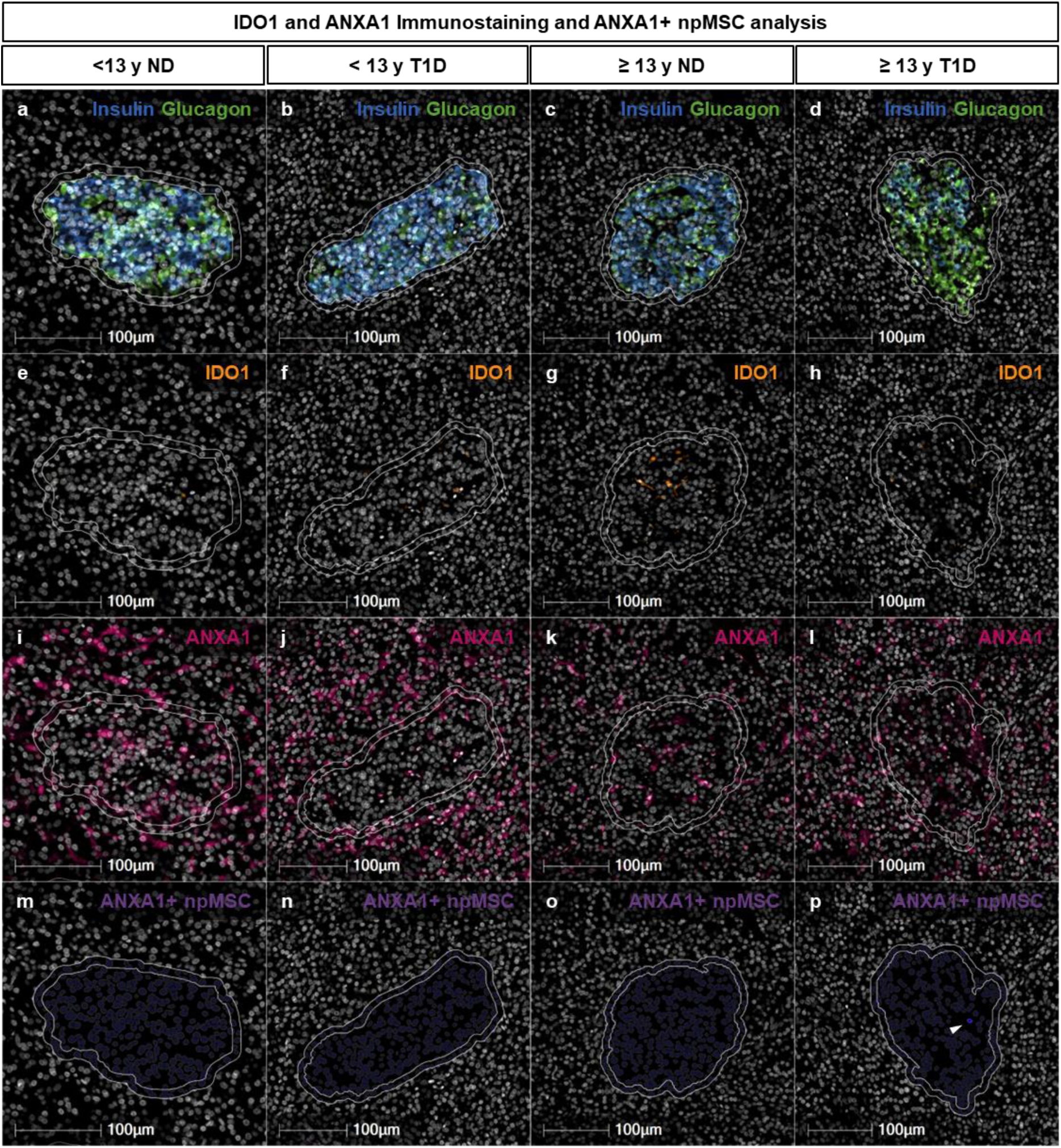
IDO1 and ANXA1 immunostaining and quantification of ANXA1+ npMSCs. Representative INS+GLU+ islets are shown. Islets were identified by insulin and glucagon immunostaining (Fig. 5a-d). IDO1 (Fig. 5e-h) and ANXA1 (Fig. 5i-l) immunostaining was conducted and ANXA1+ MSC were quantified using the HALO HighPlex FL package (npMSCs shown by white arrows, Fig. 5m-p). Each micrograph (Fig. 5a-p) shows the islet annotation (inner white annotation) and expanded annotation 10 µm from the islet periphery (outer white annotation). Donor ID: nPOD 6407 (a); nPOD 6578 (b), nPOD 6333 (c); nPOD 6362 (d). Abbreviations: < 13 y ND, < 13 years without diabetes; < 13 y T1D, < 13 years at type 1 diabetes diagnosis; ≥ 13 y ND, ≥ 13 years without diabetes; ≥ 13 y T1D, ≥ 13 years at type 1 diabetes diagnosis.

### npMSC number and density is increased at the islet periphery of insulin-containing islets in individuals with type 1 diabetes

The number and density of npMSCs within a 10 µm region immediately outside the islet (islet periphery) were quantified (Fig. 3a-d). The density of npMSCs within 10 µm of insulin-containing (INS+GLU- and INS+GLU+) but not insulin-deficient (INS-GLU+) islets was higher in individuals with type 1 diabetes compared to individuals without diabetes (Fig. 3e). Accordingly, in type 1 diabetes the density of npMSCs at the periphery of insulin-deficient islets was dramatically reduced compared to islets containing residual beta cells, consistent with an islet-protective role for npMSC. The density of npMSCs at the periphery of INS+GLU+ islets was higher in individuals who were ≥ 13 years at diagnosis of type 1 diabetes compared to those < 13 years at diagnosis (Fig. 3f). When separated by islet size (Fig. 3g-h), the more pronounced increase in npMSC number at the periphery of islets in individuals ≥ 13 years compared to < 13 years at diagnosis of diabetes was true for larger islets containing ∼64-128 or ∼265-512 cells (islet bin size 6 or 8; Fig. 3h).

In individuals without type 1 diabetes, npMSC density was also higher at the periphery of insulin-containing islets in individuals ≥ 13 years compared to individuals < 13 years (Fig. 3f). Nonetheless, the density of npMSCs at the periphery of insulin-containing islets was increased to a greater extent in individuals with type 1 diabetes irrespective of the age at diagnosis when compared to age-matched individuals without diabetes (Fig. 3f). Irrespective of age or type 1 diabetes diagnosis, the average number of npMSCs at the periphery of islets increased with islet size for insulin-containing (two-way ANOVA main effect Islet Bin Size, *p* < 0.001 for both) but not insulin-deficient (two-way ANOVA main effect Islet Bin Size, *p* = 0.61) islets (Fig. 3g-i). Thus, the number of npMSCs at the islet periphery increased with increasing numbers of beta cells.

### Intraislet npMSC density is increased within insulin-containing islets in individuals with type 1 diabetes

In accordance with our quantification of npMSCs at the islet periphery (Fig. 3a-f), the density of intraislet npMSCs within insulin-containing but not insulin-deficient islets was higher in individuals with type 1 diabetes compared to individuals without diabetes (Fig. 4a-f). However, there did not appear to be a more pronounced intraislet npMSC density in individuals ≥ 13 years at diagnosis of type 1 diabetes compared to < 13 years at diagnosis when all islet sizes were grouped together. When separated by islet size (Fig. 4g-i), only the largest INS+GLU+ islets showed increased intraislet npMSC number in individuals ≥ 13 years compared to < 13 years at diagnosis of type 1 diabetes. In agreement with our observations for npMSC number at the islet periphery, the average number of intraislet npMSCs increased with islet size for insulin-containing islets (two-way ANOVA main effect Islet Bin Size, *p* < 0.001 for both).

### npMSCs express islet-protective factors including ANXA1

Isolated “exogenous” MSCs secrete an array of cytoprotective, immunomodulatory and regenerative molecules in response to specific cues within their microenvironment to influence both islet cells and a wide range of immune cell subsets. In the current report we have determined whether the previously defined islet-protective and immunomodulatory factors, ANXA1 and IDO1, are expressed in npMSCs in the pancreas of individuals with and without type 1 diabetes (Fig. 5).

A total of 327 IDO1^+^ MSCs were identified across all pancreas sections (total pancreas including acinar, islets, ducts and vessels). The percentage of npMSCs that expressed IDO1 across all 38 individuals with and without type 1 diabetes was 1.02 ± 0.34 %. The percentage of npMSCs that were IDO1^+^ was similar for individuals with or without type 1 diabetes, irrespective of age (< 13 years at type 1 diabetes diagnosis: 2.06 ± 1.22 %; < 13 years without diabetes: 0.84 ± 0.54 %; ≥ 13 years at type 1 diabetes diagnosis: 1.15 ± 0.64 %; ≥ 13 years without diabetes: 0.25 ± 0.13 %; One-way ANOVA, F(3, 34) = 1.156, *p* = 0.32). The small number of npMSCs expressing IDO1 indicates that IDO1 is not the primary islet-protective mechanism employed by npMSCs. Further analysis was not conducted on IDO1^+^ npMSCs due to the low number of cells detected.

In contrast, a total of 16,825 ANXA1^+^ npMSCs were identified across all pancreas sections. The percentage of total pancreas npMSCs that expressed ANXA1 across all 38 individuals with and without type 1 diabetes was 33.2 % and was similar across groups, irrespective of age or type 1 diabetes diagnosis (< 13 years at type 1 diabetes diagnosis: 35.78 ± 6.20 %; < 13 years without diabetes: 27.38 ± 5.08 %; ≥ 13 years at type 1 diabetes diagnosis: 35.67 ± 5.80 %; ≥ 13 years without diabetes: 33.02 ± 5.32 %; One-way ANOVA, F(3, 34) = 0.1867, *p* = 0.73). Of the total pancreas ANXA1^+^ npMSCs identified, 7.45 % were associated with islets. Specifically, we detected 663 (3.94 %) ANXA1^+^ intraislet npMSCs and 590 (3.51 %) ANXA1^+^ npMSCs at the islet periphery (within 10 μm).

### The number and density of npMSCs expressing islet-protective ANXA1 at the islet periphery is higher in individuals with type 1 diabetes

For individuals with type 1 diabetes, the density of ANXA1^+^ npMSCs was increased at the periphery of insulin-containing but not insulin-deficient islets (Fig. 6a). When segregated according to age at type 1 diabetes diagnosis, the density of ANXA1^+^ npMSCs was enhanced at the periphery (within 10 µm) of INS+GLU+ islets for individuals ≥ 13 years, compared to individuals < 13 years at diagnosis (Fig. 6b). When further separated according to islet size (Fig. 6c-e), enhanced ANXA1+ npMSC number at the islet periphery of INS+GLU+ islets was apparent for the largest islets (islet bin size 8) containing over 256 cells (Fig. 6d). The number of ANXA1+ npMSCs at the islet periphery increased with islet size for insulin-containing islets (two-way ANOVA main effect Islet Bin Size, *p* ≤ 0.002 for both). In agreement with total npMSCs (both ANXA1^+^ and ANXA1^-^ npMSCs), this may suggest increased migration of ANXA1^+^ npMSCs to islets with an increasing number of beta cells.

**Fig. 6.**
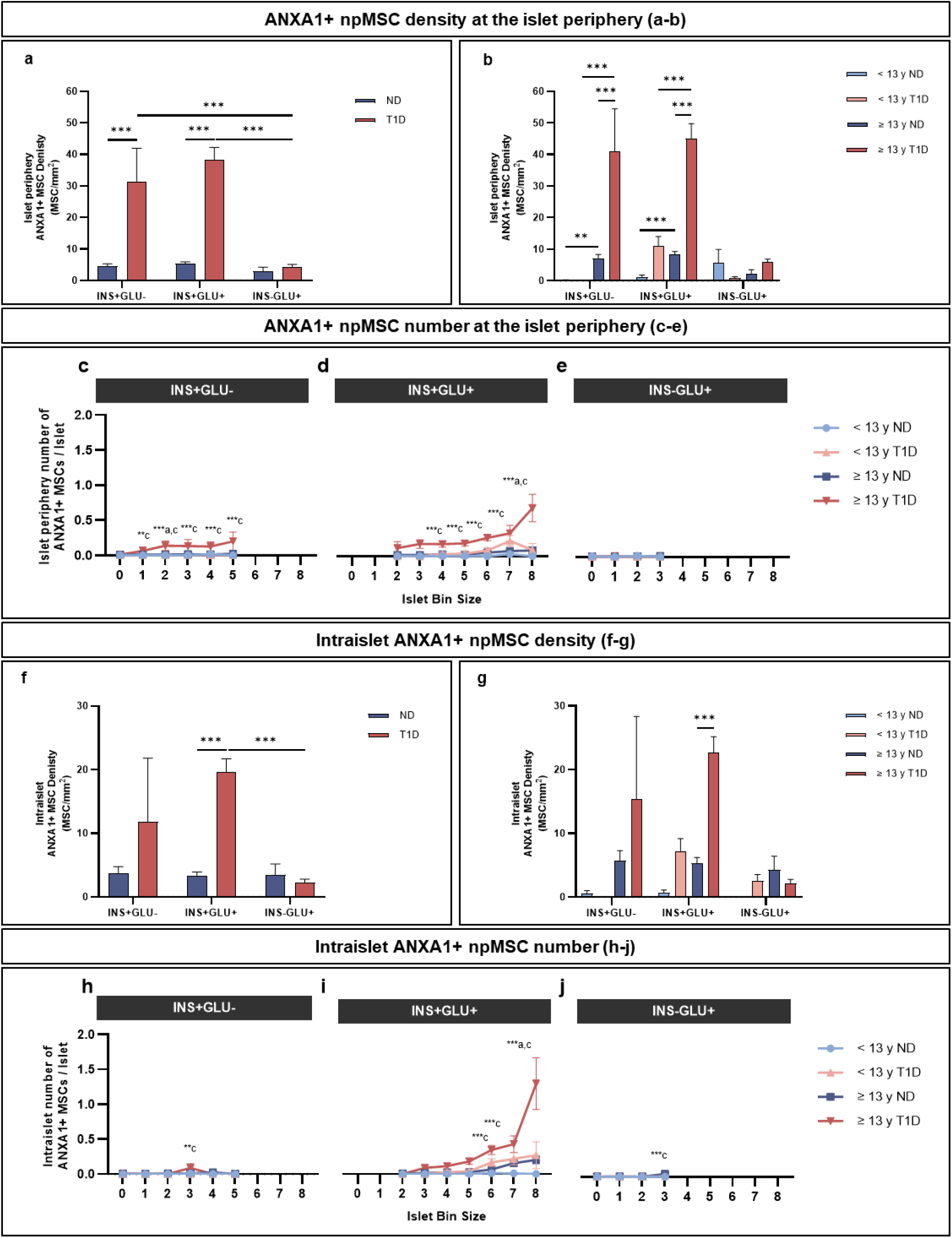
Number and density of npMSCs expressing islet-protective ANXA1 that are associated with islets. Fig. 6a-b and f-g: data derived from 26,376 individual islets from 38 individuals (*n* = 8 individuals < 13 years at type 1 diabetes diagnosis and *n* = 11 individuals ≥ 13 years at type 1 diabetes diagnosis (*n* = 19 individuals with type 1 diabetes); *n* = 8 individuals < 13 years without diabetes and *n* = 11 individuals > 13 years without diabetes (*n* = 19 individuals without diabetes)). Fig. 6c-e and h-j: islet bin sizes containing < 5 islets or bin sizes devoid of data for one or more group were excluded; data derived from *n* = 23,290 individual islets. Bars represent mean ± SEM. Two-way ordinary ANOVA with post hoc tests adjusted for multiple comparisons with Bonferroni correction. Post hoc tests in Fig. 6a and f: islets of the same endocrine cell composition; diabetes or no diabetes across islets of different endocrine cell compositions. Post hoc tests in Fig. 6b and g: type 1 diabetes and age-matched individuals without diabetes; type 1 diabetes age of diagnosis; individuals without diabetes. Comparisons in Fig. 6a-b and f-g shown with a horizontal black line between the two compared groups. Post-hoc comparisons within each islet bin size in Fig. 6c-g and h-j: < 13 years at type 1 diabetes diagnosis vs ≥ 13 years at type 1 diabetes diagnosis (in-figure label ‘a’); < 13 years at type 1 diabetes diagnosis vs < 13 years without diabetes (in-figure label ‘b’); ≥ 13 years at type 1 diabetes diagnosis vs ≥ 13 years without diabetes (in-figure label ‘c’); < 13 years without diabetes vs ≥ 13 years without diabetes (in-figure label ‘d)’. \*\*\**p* ≤ 0.001; \*\**p* ≤ 0.01. Abbreviations: < 13 y ND, < 13 years without diabetes; < 13 y T1D, < 13 years at type 1 diabetes diagnosis; ≥ 13 y ND, ≥ 13 years without diabetes; ≥ 13 y T1D, ≥ 13 years at type 1 diabetes diagnosis.

The density and number of intraislet npMSCs that express ANXA1 (Fig. 6f-j) mostly aligned with the data for total (both ANXA1^+^ and ANXA1^-^ npMSCs) intraislet npMSCs within INS+GLU+ islets. An exception to this was that intraislet ANXA1^+^ npMSC density between individuals < 13 years with or without type 1 diabetes was similar.

### Whole pancreas and individual islet immune cell infiltration

Total pancreas CD45^+^ cell number was quantified as an index of inflammation. As expected, when categorised by age and diabetes status, whole pancreas inflammation was increased in individuals with type 1 diabetes compared to individuals without (two-way ANOVA main effect diabetes, *p* < 0.001). When separated by age, irrespective of diabetes diagnosis, total pancreas inflammation was similar (*p* = 0.76).

An islet was characterised as inflamed if there were more than five associated immune cells. This was determined by quantifying the total number of CD45^+^ cells located either inside the islet or within 10 µm of the islet periphery (Fig. 7a-d) [33, 34]. Very few (0.15%) INS+GLU+ islets from individuals without diabetes were categorised as inflamed, as expected [34]. Therefore, at the individual islet level, inflammation was characterised further for INS+GLU+ islets from individuals with type 1 diabetes only. The proportion of INS+GLU+ islets that were inflamed was 53.6% (*n* = 126 islets) and 13.6% (*n* = 130 islets) for individuals < 13 years or ≥ 13 years at diagnosis of type 1 diabetes, respectively. The average number of immune cells associated with each inflamed islet was higher for individuals < 13 years at diagnosis of type 1 diabetes, compared to ≥ 13 years at diagnosis of type 1 diabetes (mean ± SEM; 27.48 ± 3.35 vs 15.84 ± 1.82 CD45^+^ cells; Unpaired t test; t(254) = 3.078, *p* = 0.002). This is consistent with the increased insulitis and more aggressive destruction of beta cells reported to occur in individuals with T1DE1 compared to T1DE2 [29, 30]. Whilst the number of beta cells per INS+GLU+ islet was not different between age groups, inflamed islets contained more beta cells than islets not under obvious immune attack (Fig. 7e), suggesting that larger islets may be preferentially infiltrated by immune cells.

**Fig. 7.**
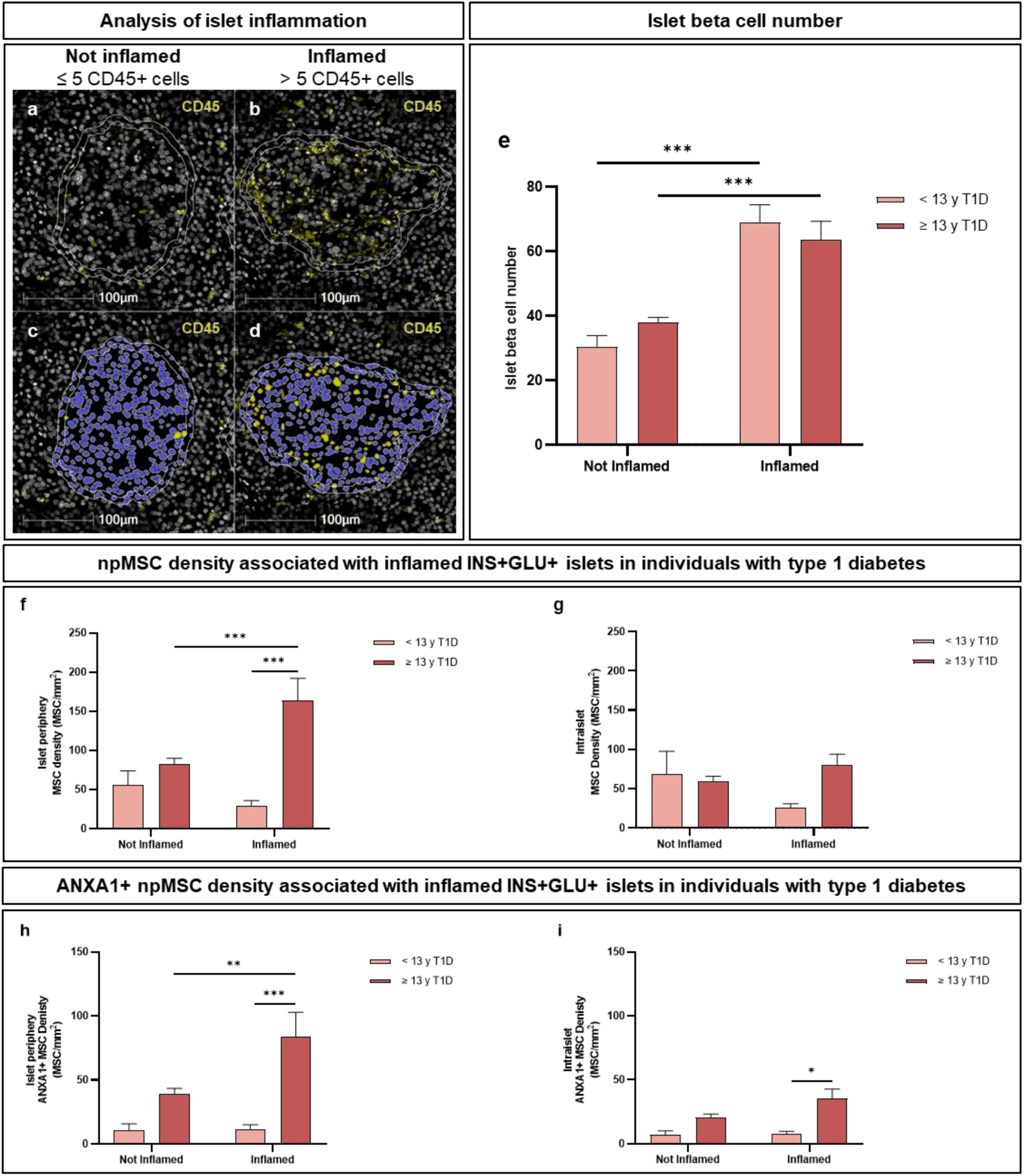
npMSCs are preferentially associated with inflamed islets in individuals older than 13 years at type 1 diabetes diagnosis. The number of CD45+ cells associated with islets (that were either within 10 µm of the islet periphery or inside the islet; Fig. 7a-b) were quantified and, islets were classified as not inflamed (≤ 5 CD45+ cells; Fig. 7c) or inflamed (> 5 CD45+ cells; Fig. 7d). Islets shown in Fig. 7a-d as inner white annotation and islet annotations expanded 10 µm shown by the outer annotation. Fig. 7e-i: data were derived from 1191 individual islets from *n* = 8 individuals < 13 years at type 1 diabetes diagnosis and *n* = 11 individuals ≥ 13 years at type 1 diabetes diagnosis. Bars represent mean ± SEM. Two-way ordinary ANOVA with post hoc tests adjusted for multiple comparisons with Bonferroni correction. Post hoc tests in Fig. 7e-i: between type 1 diabetes age of diagnosis at the same level of inflammation; between inflammation status for the same age of diagnosis. Comparisons shown with a horizontal black line between the two compared groups. Donor ID: 6578, nPOD (a-d). \*\*\**p* ≤ 0.001; \*\**p* ≤ 0.01; \**p* ≤ 0.05. Abbreviations: < 13 y T1D, < 13 years at type 1 diabetes diagnosis; ≥ 13 y T1D, ≥ 13 years at type 1 diabetes diagnosis.

npMSC density (ANXA1^+^ and ANXA1^-^) at the periphery, but not within (intraislet), inflamed INS+GLU+ islets was increased in individuals ≥ 13 years at diagnosis of type 1 diabetes compared to islets that were not inflamed from the same group and compared to inflamed islets in individuals < 13 years at diagnosis (Fig. 7f-g). The density of npMSCs that express ANXA1 at the periphery of inflamed INS+GLU+ islets displayed similar trends to those of total npMSCs (ANXA1^+^ and ANXA1^-^ (Fig. 7h)). Intraislet ANXA1^+^ npMSC density was increased in inflamed INS+GLU+ islets of individuals ≥ 13 years compared to < 13 years at type 1 diabetes diagnosis (Fig. 7i).

### Viability and proliferation of cytokine-exposed human MSCs in vitro

We investigated the ability of MSCs to survive and proliferate under cytokine exposures of increasing “aggression” and duration (Fig. 8a-d). MSC viability was reduced in response to the more aggressive cytokine cocktail (IFN-γ + IL-1β + TNF-α), compared to exposure to less aggressive conditions (IFN-α alone or IFN-γ + IL-1β) or to the no cytokine control following three- or seven-day exposure (Fig. 8a-b). The percentage of MSCs that had divided over the course of the three-day experiment was also reduced in response to the more aggressive cytokine cocktail compared to the no cytokine control (Fig. 8c).

**Fig. 8.**
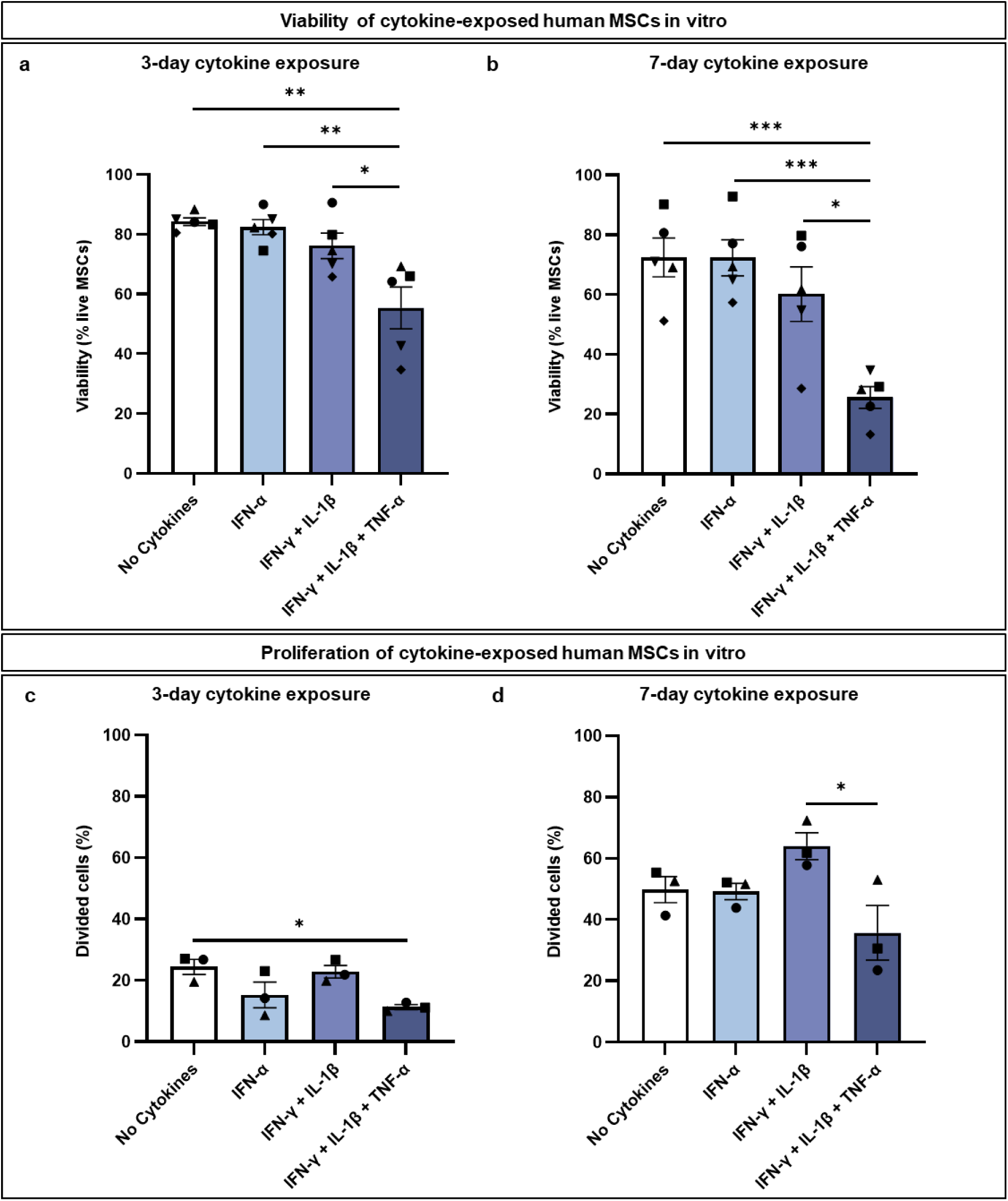
Viability and proliferation of cytokine-exposed human MSCs in vitro. Human MSCs were exposed to a cytokine condition of increasing aggression and viability was assessed after three (a) or seven (b) days. The percentage of divided cells, as calculated by the FlowJo proliferation algorithm, was used to determine proliferation of CellTrace-stained cells after three (c) or seven (d) days. Each symbol represents an independent experiment (*n* = 5 for viability and *n* = 3 for proliferation). Bars represent mean ± SEM. Human MSCs between passages 5-8 were used for all experiments. One-way ANOVA with post hoc tests adjusted for multiple comparisons with Bonferroni corrections. Comparisons in Fig. 8a-d shown with a horizontal black line between the two compared groups. \*\*\**p* ≤ 0.001; \*\**p* ≤ 0.01; \**p* ≤ 0.05.

Overall, exposure of MSCs to an aggressive cytokine combination led to increased cell death and reduced proliferation. This is particularly reflective of the cytokine exposure expected around islets in individuals < 13 years at diagnosis of type 1 diabetes, where immune cell infiltration is more intense [29, 30]. This may indicate that npMSCs are less able to survive in situ at the islet periphery in individuals < 13 years at diagnosis of type 1 diabetes.

## Discussion

To our knowledge, this is the first report providing a comprehensive immunohistological characterisation of npMSCs in individuals with and without type 1 diabetes. Using human pancreas tissue sections from both the nPOD and EADB collections, we have been able to monitor npMSC number, density and islet-protective phenotype close to diabetes onset (within 2 years) and to explore their association with inflamed islets in individuals with type 1 diabetes.

The current study introduces a methodological pipeline for phenotyping npMSCs on an individual-islet basis across whole-slide scans using the quantitative digital pathology platform, HALO. This pipeline provides a reproducible framework for analysing cell phenotypes in individual regions of interest in a high-throughput manner across entire tissue sections. This pipeline can be adapted to identify other cell phenotypes across a range of tissue types, highlighting its applicability to different biological and disease contexts.

In vivo, native MSCs characteristically display similar phenotypic features to those reported in isolated (“exogenous”), culture-expanded MSCs. MSCs have markers in common with other microvascular and stromal cell types including, but not limited to, endothelial cells [35], pericytes [24, 36, 37] and stellate cells [22, 38, 39]. As such no single phenotypic marker distinguishes MSCs from other cell types. Thus, we have utilised multiplex fluorescent immunohistochemistry, using Opal 6-colour methodology, incorporating a comprehensive panel of human MSC markers (CD73^+^, CD90^+^, CD105^+^, CD31^-^, CD34^-^, CD45^-^) to unequivocally define MSC populations in human pancreas tissue from individuals with and without type 1 diabetes.

Automated quantitative image analysis following whole-slide scanning and multispectral unmixing, has facilitated high throughput analysis of 53,375 npMSCs across 38 individuals with and without type 1 diabetes. We have identified MSC populations in the exocrine tissue as well as intraislet MSCs, in accordance with reports that MSCs can be isolated from pancreatic ductal epithelial cells [40], as well as acinar and endocrine pancreas tissue [20–26, 41, 42]. Moreover, we have demonstrated significant alterations in npMSC number and density in individuals with type 1 diabetes.

MSCs respond to specific cues within their microenvironment and migrate to sites of injury, including pancreatic islets [19], where they exert their immunomodulatory and regenerative functions. Our detailed histological studies of the human pancreas have demonstrated that the number and density of exocrine npMSCs within the immediate vicinity (10 µm) of islets (specifically those still containing beta cells) is increased in individuals with type 1 diabetes. Furthermore, the percentage of INS+GLU+ islets which have npMSCs localised to the islet periphery is also markedly increased in type 1 diabetes compared to individuals without type 1 diabetes. Our analysis has intentionally focused on pancreas samples from individuals with recent-onset type 1 diabetes (≤ 2 years), where islet inflammation and immune attack is still ongoing [34].

This has enabled us to further demonstrate that npMSCs are preferentially associated with inflamed islets (more than five CD45^+^ immune cells either within the islet or within 10 µm of the islet periphery), indicating the selective homing of MSCs to inflamed human pancreatic islets. Accordingly, whilst we detected alterations in the number and density of npMSCs in close proximity to pancreatic islets, npMSC number and density across total pancreas area was similar between individuals with and without type 1 diabetes.

We have further demonstrated that, in type 1 diabetes, npMSCs are preferentially localised to larger islets (bin size ≥ 6) containing a higher number of residual beta cells. This is perhaps not surprising given the established homing capacity of MSCs [19] towards islet-expressed chemokines including CXCL12 and CXC3L1 [19, 43]. Thus, it is expected that islets with a higher number of residual beta cells would synthesise and release more MSC chemoattractants generating a stronger chemokine gradient for MSC migration.

Whilst our data support the notion that MSCs home to inflamed islets in the human pancreas during the progression of type 1 diabetes, it is beyond the scope of this study to definitively define the tissue origin of “npMSCs” identified in the pancreas. Thus, although, we have standardised our terminology to npMSC for all MSC populations detected in the exocrine and endocrine pancreas, we acknowledge that npMSCs may well represent a population of ‘homing’ MSCs that originated from other tissue sources, such as bone marrow, as previously reported [19]. It is also plausible that there are subpopulations of npMSCS that have been generated through epithelial to mesenchymal transition from acinar or pancreatic ductal cells [44, 45].

One compelling mechanism to explain the increased number of npMSCs in type 1 diabetes is that subpopulation(s) of MSCs within the exocrine and/or endocrine pancreas arise due to the differentiation of pericytes into MSCs/pericyte-MSCs [46, 47]. Pericytes wrap around endothelial cells, behave as stem cells and can give rise to progenitor cells and MSCs when removed from the body and expanded ex vivo [46]. This process may reflect changes that occur in vivo under inflammatory conditions, such as those seen during type 1 diabetes pathogenesis and progression. Pericytes are known to become myofibroblasts in fibrotic diseases, including islet vascular fibrosis [41] and to form native MSCs/pericyte-MSCs to mediate tissue repair [37].

Indeed, pro-inflammatory cytokines influence pericytes to differentiate into native MSCs by loosening their physical association with endothelial cells and inducing their proliferation, migration and secretion of immunomodulatory factors [46, 48]. Whilst islet pericyte density has been reported to remain unchanged in type 1 diabetes, islet pericyte capillary coverage is reduced [49]. This may suggest that pericytes detach from the vasculature and differentiate into reparative npMSCs in type 1 diabetes. This hypothesis is supported by the current study which demonstrates an increased density of npMSCs associated with insulin-containing islets in individuals with type 1 diabetes.

Our studies consistently demonstrate that the increased number of npMSCs in individuals with type 1 diabetes is more pronounced in individuals with an older age at diagnosis (≥ 13 years). This could be explained by differences in the migratory capacity of MSCs with age. However, previous studies have demonstrated that MSC migration is reduced rather than enhanced with increasing age [50]. Alternatively, residual beta cells in the pancreas of individuals with later onset type 1 diabetes may be better able to attract MSCs due to differences in their chemokine expression profile. Our in vitro measurements of MSC viability following exposure to pro-inflammatory cytokines suggest that the differences in npMSC number observed in situ are at least partially attributable to their ability to survive differentially aggressive cytokine cocktails with varying age of diabetes onset. Thus, our in vitro observations indicate that the increased number of npMSCs in individuals diagnosed with type 1 diabetes after the age of 13 is likely to be related to the ability of MSCs to survive the less aggressive immune cell infiltration and attack typically associated with later diagnosis (≥ 13 years) of type 1 diabetes [29, 30]. Accordingly, whilst we have shown that MSC viability in vitro is not influenced by a 7-day exposure to IFN-α alone or a dual combination of IFN-γ and IL-1β, our studies have consistently demonstrated a dramatic reduction in MSC viability following exposure to a more aggressive cocktail of IFN-γ, IL-1β and TNF-α.

In health, the density of npMSCs either within or in close proximity to insulin-containing islets increased with age, albeit to a lesser extent than that seen with type 1 diabetes. Notably, whole pancreas (total pancreas area including acinar, islets, ducts and vessels) MSC and immune cell (CD45^+^ cells) density remained similar across age groups, suggesting that the observed increase in npMSC density associated with islets is unlikely to reflect age-related increases in inflammation. This is consistent with the relatively young age of the individuals included in this study (all individuals ≤ 27 years), where age-dependent pancreas inflammation would still be minimal [34]. These differences may be explained by the expected increase in beta cell maturation and function across age groups which is suggested to occur over the first three decades of life [51–53]. Indeed, the proportion of beta cells expressing endoplasmic reticulum stress, insulin secretion or autophagy-related genes was reported to increase in healthy donors 10-19 years compared to < 10 years [53]. Therefore, it is plausible that insulin-containing islets from healthy individuals ≥ 13 years in the present study produced increased levels of chemokines related to insulin secretion and tissue repair and provided a greater stimulus for MSC migration than in individuals < 13 years.

Previous studies have revealed therapeutic mechanisms by which MSCs isolated from various post-natal tissues (including bone marrow, adipose, kidney and pancreas) then subsequently “culture-expanded” improve islet function and survival [12, 54–59], as well as immune cell activity [5, 60, 61]. MSCs secrete an array of cytoprotective, immunomodulatory and regenerative molecules [7, 62–64] in response to specific cues within their microenvironment to influence both islet cells [54, 55, 65] and a wide range of immune cell subsets.

ANXA1 is an MSC-secreted islet G-protein coupled receptor ligand and is a key modulator of MSC-mediated improvements in islet functional survival following cytokine exposure in vitro [54, 55, 65]. In the current study we have demonstrated that a significant proportion (> 25 percent for individuals with or without type 1 diabetes) of npMSCs within the immediate vicinity of human pancreatic islets express ANXA1 protein. Whilst the proportion of npMSCs at the islet periphery immunopositive for ANXA1 was comparable for individuals with and without type 1 diabetes, the number and density of ANXA1 immunopositive npMSCs was increased in individuals with type 1 diabetes. This reflects the diabetes-related increase in total (irrespective of ANXA1 expression) npMSC number and density at the islet periphery. Thus, diabetes-related alterations in the number and density of ANXA1 immunopositive npMSCs do not appear to be due to the induction of ANXA1 expression in npMSCs exposed to diabetogenic stressors. Accordingly, isolated and culture-expanded MSCs constitutively express high levels of ANXA1 in the absence of cytokine exposure [54, 55].

Immunosuppression by human MSCs is known to be mediated by IDO1, an enzyme that catalyses the rate-limiting step in the degradation of tryptophan, along the kynurenine pathway [66]. Thus, MSCs down regulate T cell proliferation and functioning, in part via IDO-mediated tryptophan catabolism [5, 66, 67]. We therefore sought to determine whether npMSCs in the human pancreas of individuals with recent onset type 1 diabetes, where immune attack is ongoing, express IDO1. In contrast to the relatively high proportion of npMSCs (> 25 percent) found to express the anti-inflammatory protein, ANXA1, our study shows that less than two percent of npMSCs are immunopositive for IDO1. The current observations therefore indicate that a protective role of npMSCs in the human pancreas is likely to be at least in part attributable to the anti-inflammatory functions of ANXA1 [68, 69], and that IDO1 is not a key therapeutic mechanism through which npMSCs modulate immune cells infiltrating the islets.

MSCs have a plethora of anti-inflammatory, immunomodulatory and regenerative properties (reviewed in [60, 70]) including the functional capacity to reduce CD4^+^, CD8^+^ T cell, as well as B cell proliferation and activity. Our in-situ studies of the human pancreas suggest that MSCs are recruited to islets undergoing immune attack where they express anti-inflammatory islet-protective factors, including ANXA1, to preserve endogenous islet beta cell survival. Accordingly, in type 1 diabetes both the density of total npMSCs and of ANXA1^+^ npMSCs at the periphery of insulin deficient islets is dramatically reduced compared to islets containing residual beta cells. Furthermore, the density and number of ANXA1^+^ npMSCs observed at the periphery of insulin containing islets is higher in individuals with later onset type 1 diabetes (≥ 13 years), corresponding to the less aggressive rate of type 1 diabetes progression known to occur in these individuals [29, 30]. Thus, our current study supports an islet-protective role for endogenous npMSCs in delaying type 1 diabetes progression.

Our conclusions are based on data generated from individuals of European decent. Future studies are required to confirm whether the number, distribution or immunomodulatory phenotype of npMSCs differs with ethnicity, which may be expected due to the heterogeneity of type 1 diabetes between ethnic groups (reviewed in [31]). Our study intentionally focused on individuals with short-duration type 1 diabetes, where immune cell infiltration is still ongoing [34]. This enabled us to demonstrate that npMSCs were preferentially associated with inflamed islets in individuals ≥ 13 years at diagnosis of type 1 diabetes. Our future studies will be directed towards investigating whether the increased npMSC density associated with individuals ≥ 13 years is maintained into longer duration disease where beta cell loss is more pronounced, and islet inflammation is relatively resolved.

Altogether, the results presented here suggest a protective role for npMSCs in delaying the progression of type 1 diabetes. We suggest that npMSCs localised at the periphery of insulin containing islets may play a protective role to limit CD8^+^ T cell-mediated destruction of islets. In accord with this, our detailed observations demonstrate that; (1) npMSCs are more abundant in the vicinity of insulin-containing islets when compared to insulin-deficient islets; (2) npMSCs are more abundant in individuals with older onset type 1 diabetes (age at diagnosis ≥ 13 years), corresponding with a less aggressive immune cell infiltration; (3) npMSCs are preferentially associated with larger islets containing a greater number of residual beta cells; and (4) npMSCs express therapeutic factors, namely ANXA1, that have established roles in protecting islets from cytokine induced apoptosis and preserving insulin secretory function [54, 55]

Therapeutic strategies to modulate the homing of endogenous or systemically infused exogenous MSCs to pancreatic islets represent a promising way to enhance the clinical applications of MSCs to preserve endogenous beta cells. In addition, enhancing the resilience of MSCs to aggressive cytokine exposure, may improve their clinical application in individuals with younger onset type 1 diabetes where MSC survival is challenged by the aggressive inflammatory pancreas environment. Overall, our data suggest that therapeutically administered MSCs are likely to be more effective at preserving endogenous beta cell mass in individuals with later onset type 1 diabetes where immune cell infiltration is less aggressive and MSCs have a greater ability to survive and exert their reparative functions.

## Supporting information

Electronic Supplementary Material

## Acknowledgements

The authors would like to thank Dr Katy Murrall for providing calculations for the determination of islet bin sizes within this manuscript. This research was performed with the support of the Network for Pancreatic Organ donors with Diabetes (nPOD; RRID:SCR_014641), a collaborative type 1 diabetes research project supported by Breakthrough T1D and The Leona M. & Harry B. Helmsley Charitable Trust (Grant 3-SRA-2023-1417-S-B). The content and views expressed are the responsibility of the authors and do not necessarily reflect the official view of nPOD. Organ Procurement Organizations (OPO) partnering with nPOD to provide research resources are listed at https://npod.org/for-partners/npod-partners/. The authors gratefully acknowledge the Exeter Centre for Cytomics at the University of Exeter for their support and assistance with this work. This study has been delivered through the National Institute for Health and Care Research (NIHR) Exeter Biomedical Research Centre (BRC). The views expressed are those of the authors and not necessarily reflect those of Breakthrough T1D, Diabetes UK, the NIHR or the Department of Health and Social Care.

## Data availability

All data are available in the main text or the electronic supplementary material (ESM). They are available from the corresponding authors upon reasonable request. For the purpose of open access, the author has applied a Creative Commons Attribution (CC BY) licence to any Author Accepted Manuscript version arising from this submission.

## Authors relationships and activities

The authors declare that there are no relationships or activities that might bias, or be perceived to bias, their work.

## Contribution statement

RDT: Conception and design, financial support, acquisition, analysis and interpretation of data, drafting and critical review of the manuscript. JA: Acquisition of data and critical review of the manuscript. NM: Conception and design, financial support, interpretation of data, critical review of manuscript. ME: Conception and design, analysis and interpretation of data, critical review of manuscript. SR: Conception and design, financial support, interpretation of data, critical review of manuscript. CR: Conception and design, financial support, interpretation of data, drafting and critical review of the manuscript. All authors approved the final version of the manuscript. CR was principal investigator supervising this study and is the guarantor for this work.

## Abbreviations

ANXA1: Annexin A1
EADB: Exeter Archival Diabetes Biobank
GLU: Glucagon
HIER: Heat induced epitope retrieval
IDO1: Indoleamine 2,3-dioxygenase 1
INS: Insulin
MSC: Mesenchymal stromal cell
nPOD: Network for pancreatic organ donors with diabetes
npMSC: Native human pancreatic mesenchymal stromal cell
T1DE1: Type 1 diabetes endotype 1
T1DE2: Type 1 diabetes endotype 2

