## Supplementary material for "Characterisation of native human pancreatic mesenchymal stromal cells in type 1 diabetes": Electronic Supplementary Material

**Content:**

ESM Tables

ESM Figures

### ESM Tables

**ESM Table 1** Clinical characteristics of pancreas donors included in this study from the EADB and nPOD collections.

| Tissue Collection | Donor ID RRID | Group | Age (years) | Sex | T1D duration (y) | BMI (kg/m$^2$) | C-Peptide (ng/mL) | HbA1C | Islet Autoantibody Positivity |
|---|---|---|---|---|---|---|---|---|---|
| EADB | E428 SAMN46311904 | < 13 y T1D | 5 | M | 0.02 | Unknown | Unknown | Unknown | Unknown |
| EADB | E375 SAMN46311885 | < 13 y T1D | 11 | F | 0.02 | Unknown | Unknown | Unknown | Unknown |
| EADB | SC115 SAMN46311955 | < 13 y T1D | 1 | F | 0.01 | Unknown | Unknown | Unknown | Unknown |
| EADB | 485/88 SAMN46312114 | < 13 y ND | 2 | F | N/A | Unknown | Unknown | Unknown | Unknown |
| EADB | 12426 SAMN46312017 | < 13 y ND | 5 | Unknown | N/A | Unknown | Unknown | Unknown | Unknown |
| EADB | 315/89 SAMN46312086 | < 13 y ND | 9 | M | N/A | Unknown | Unknown | Unknown | Unknown |
| EADB | E386 SAMN46311889 | ≥ 13 y T1D | 15 | M | 0.5 | Unknown | Unknown | Unknown | Unknown |
| EADB | SC57 SAMN46311975 | ≥ 13 y T1D | 18 | F | 0.02 | Unknown | Unknown | Unknown | Unknown |
| EADB | SC76 SAMN46311979 | ≥ 13 y T1D | 20 | M | 0.06 | Unknown | Unknown | Unknown | Unknown |
| EADB | 146/66 SAMN46312039 | ≥ 13 y ND | 18 | F | N/A | Unknown | Unknown | Unknown | Unknown |
| EADB | PAN8 SAMN46312163 | ≥ 13 y ND | 19 | Unknown | N/A | Unknown | Unknown | Unknown | Unknown |
| EADB | 333/66 SAMN46312091 | ≥ 13 y ND | 16 | M | N/A | Unknown | Unknown | Unknown | Unknown |

| | | | | | | | | | |
|---|---|---|---|---|---|---|---|---|---|
| nPOD | 6533 SAMN18242777 | < 13 y T1D | 3.75 | F | 0 | 17.7 | 0.17 | 11.4 | IA2A+ mIAA+ ZnT8A+ |
| nPOD | 6534 SAMN18242778 | < 13 y T1D | 4.19 | F | 0 | 22.6 | 0.05 | 15.6 | IA-2A+ |
| nPOD | 6578 SAMN33284295 | < 13 y T1D | 11.95 | F | 0 | 22.5 | 0.35 | 13.6 | IA2A+ ZnT8A+ |
| nPOD | 6209 SAMN15879265 | < 13 y T1D | 5 | F | 0.25 | 15.9 | 0.1 | Unknown | IA-2A+ ZnT8A+ mIAA+ |
| nPOD | 6371 SAMN15879424 | < 13 y T1D | 12.5 | F | 2 | 16.6 | 0.11 | 9.5 | GADA+ IA-2A+ mIAA+ ZnT8A+ |
| nPOD | 6407 SAMN15879460 | < 13 y ND | 4.6 | F | N/A | 16 | 5.35 | 5.5 | Negative |
| nPOD | 6488 SAMN15879541 | < 13 y ND | 4.6 | F | N/A | 16.8 | 8.67 | 5.6 | Negative |
| nPOD | 6382 SAMN15879435 | < 13 y ND | 4.7 | F | N/A | 17.2 | 5.21 | 5.3 | Negative |
| nPOD | 6293 SAMN15879347 | < 13 y ND | 9 | F | N/A | 18.6 | 2.22 | Unknown | Negative |
| nPOD | 6413 SAMN15879466 | < 13 y ND | 10.1 | F | N/A | 19 | 5.27 | 5.6 | Negative |
| nPOD | 6228 SAMN15879284 | ≥ 13 y T1D | 13 | M | 0 | 17.4 | 0.1 | 13.3 | GADA+ IA-2A+ ZnT8A+ |
| nPOD | 6563 SAMN30386851 | ≥ 13 y T1D | 14.56 | F | 0 | 25.5 | 1.04 | 9.6 | IA2A+ |
| nPOD | 6551 SAMN25652262 | ≥ 13 y T1D | 20.7 | M | 0.58 | 23.1 | 0.11 | 6.4 | GADA+ IA-2A+ mIAA+ ZnT8A+ |
| nPOD | 6520 SAMN18053203 | ≥ 13 y T1D | 21.61 | M | 0 | 29.3 | 0.37 | 11.9 | GADA+ IA-2A+ ZnT8A+ |
| nPOD | 6362 SAMN15879415 | ≥ 13 y T1D | 24.9 | M | 0 | 28.5 | 0.38 | 10 | GADA+ |

| | | | | | | | | | |
|---|---|---|---|---|---|---|---|---|---|
| nPOD | 6550 SAMN25652261 | ≥ 13 y T1D | 25.06 | M | 0 | 16.4 | <0.02 | 14 | GADA+ ZnT8A+ |
| nPOD | 6579 SAMN33284296 | ≥ 13 y T1D | 13.91 | F | 1.167 | 18.4 | 0.31 | 15 | GADA+ mIAA+ |
| nPOD | 6469 SAMN15879522 | ≥ 13 y T1D | 26.06 | F | 1.5 | 26.9 | 0.66 | 7.4 | GADA+ |
| nPOD | 6501 SAMN15879554 | ≥ 13 y ND | 12.85 | M | N/A | 15.8 | 9.69 | 5.2 | Negative |
| nPOD | 6374 SAMN15879427 | ≥ 13 y ND | 14 | F | N/A | 18.9 | 13.42 | 6.1 | Negative |
| nPOD | 6548 SAMN25652259 | ≥ 13 y ND | 20.24 | M | N/A | 23.8 | 4.04 | 5.7 | Negative |
| nPOD | 6339 SAMN15879393 | ≥ 13 y ND | 23.3 | M | N/A | 25 | 10.56 | 5.3 | Negative |
| nPOD | 6431 SAMN15879484 | ≥ 13 y ND | 13.79 | M | N/A | 23.1 | 1.32 | 5.4 | Negative |
| nPOD | 6271 SAMN15879325 | ≥ 13 y ND | 17 | M | N/A | 24.4 | 11.47 | Not reported | Negative |
| nPOD | 6232 SAMN15879288 | ≥ 13 y ND | 14 | F | N/A | 20.83 | 19.5 | Not reported | Negative |
| nPOD | 6333 SAMN15879387 | ≥ 13 y ND | 27.1 | F | N/A | 24.9 | 9.37 | 4.7 | Negative |

Abbreviations: < 13 y ND, < 13 years without diabetes; ≥ 13 y ND, ≥ 13 years without diabetes; < 13 y T1D, < 13 years at type 1 diabetes diagnosis; ≥ 13 y T1D, ≥ 13 years at type 1 diabetes diagnosis.

**ESM Table 2** Optimised Opal panel to identify npMSCs.

| Antigen | Antibody (clone)<br>Product number<br>Host species<br>RRID | HIER | HRP Secondary | Opal fluorophore | Position in staining panel |
|---|---|---|---|---|---|
| CD90 | CD90 (EPR3132)<br>Ab92574; Abcam<br>Rb mAb<br>AB_10563647 | Citrate<br>pH 6 | Opal Ms/Rb<br>HRP | 570 | 1 |
| CD105 | CD105 (EPR10145-12)<br>Ab169545; Abcam<br>Rb mAb<br>AB_2894873 | TE<br>pH 9 | Opal Ms/Rb<br>HRP | 690 | 2 |
| CD31 | CD31 (EPR3094)<br>Ab76533; Abcam<br>Rb mAb<br>AB_1523298 | TE<br>pH 9 | Opal Ms/Rb<br>HRP | 480 | 3 |
| CD45 | CD45 (2B11 + PD7/26)<br>M0701; Agilent<br>Ms mAb<br>AB_2314143 | Citrate<br>pH 6 | Opal Ms/Rb<br>HRP | 620 | 4 |
| CD73 | CD73 (D7F9A)<br>13160; Cell Signaling<br>Rb mAb<br>AB_2716625 | Citrate<br>pH 6 | Opal Ms/Rb<br>HRP | 520 | 5 |
| CD34 | CD34 (QBEnd 10)<br>MA110202; Invitrogen<br>Ms mAb<br>AB_11156010 | TE<br>pH 9 | Opal Ms/Rb<br>HRP | 780 | 6 |

Abbreviations: HRP, horseradish peroxidase; mAb, monoclonal antibody; Ms, mouse; Rb, rabbit; TE, tris EDTA.

**ESM Table 3** Optimised Opal panel to identify islets and MSC-derived islet-protective factors.

| Antigen | Antibody (clone)<br>Product number<br>Host species<br>RRID | HIER | HRP Secondary | Opal fluorophore | Position in staining panel |
|---|---|---|---|---|---|
| CD90 | CD90 (EPR3132)<br>Ab92574; Abcam<br>Rb mAb<br>AB_10563647 | Citrate<br>pH 6 | Opal Ms/Rb<br>HRP | 570 | 1 |
| IDO1 | IDO1 (EPR20374)<br>Ab211017; Abcam<br>Rb mAb<br>AB_2936946 | TE<br>pH 9 | Opal Ms/Rb<br>HRP | 480 | 2 |
| ANXA1 | ANXA1 (EPR19342)<br>Ab214486; Abcam<br>Rb mAb<br>AB_2890907 | TE<br>pH 9 | Opal Ms/Rb<br>HRP | 520 | 3 |
| CD45 | CD45 (2B11 + PD7/26)<br>M0701; Agilent<br>Ms mAb<br>AB_2314143 | Citrate<br>pH 6 | Opal Ms/Rb<br>HRP | 620 | 4 |
| Insulin | Insulin (ICBTACLS)<br>14-9769-82; Invitrogen<br>Ms mAb<br>AB_2573014 | Citrate<br>pH 6 | Opal Ms/Rb<br>HRP | 690 | 5 |
| Glucagon | Glucagon (K79bB10)<br>Ab10988, Abcam<br>Ms mAb<br>AB_297642 | Citrate<br>pH 6 | Opal Ms/Rb<br>HRP | 780 | 6 |

Abbreviations: HRP, horseradish peroxidase; mAb, monoclonal antibody; Ms, mouse; Rb, rabbit; TE, tris EDTA.

**ESM Table 4** Number and percentage of islets comprising different endocrine cell compositions among individual donors with and without type 1 diabetes from the EADB and nPOD collections.

| | | Absolute number of islets (*n*) | | | | Percentage of total islets (%) | | |
|---|---|---|---|---|---|---|---|---|
| **Donor ID** | **Group** | **INS+GLU-** | **INS+GLU+** | **INS-GLU+** | **Total Islets** | **INS+GLU-** | **INS+GLU+** | **INS-GLU+** |
| E428B | < 13 y T1D | 4 | 11 | 157 | 172 | 2.33 | 6.40 | 91.28 |
| E375 | < 13 y T1D | 0 | 6 | 83 | 89 | 0.00 | 6.74 | 93.26 |
| SC115 | < 13 y T1D | 2 | 9 | 193 | 204 | 0.98 | 4.41 | 94.61 |
| 6533 | < 13 y T1D | 20 | 11 | 782 | 813 | 2.46 | 1.35 | 96.19 |
| 6534 | < 13 y T1D | 4 | 43 | 184 | 231 | 1.73 | 18.61 | 79.65 |
| 6578 | < 13 y T1D | 57 | 94 | 143 | 294 | 19.39 | 31.97 | 48.64 |
| 6209 | < 13 y T1D | 3 | 39 | 532 | 574 | 0.52 | 6.79 | 92.68 |
| 6371 | < 13 y T1D | 1 | 22 | 251 | 274 | 0.36 | 8.03 | 91.61 |
| 6407 | <13 y ND | 193 | 281 | 93 | 567 | 34.04 | 49.56 | 16.40 |
| 6488 | <13 y ND | 937 | 553 | 33 | 1523 | 61.52 | 36.31 | 2.17 |
| 6382 | <13 y ND | 443 | 403 | 47 | 893 | 49.61 | 45.13 | 5.26 |
| 6293 | <13 y ND | 106 | 549 | 76 | 731 | 14.50 | 75.10 | 10.40 |
| 6413 | <13 y ND | 247 | 304 | 28 | 579 | 42.66 | 52.50 | 4.84 |
| 12426 | <13 y ND | 360 | 498 | 20 | 878 | 41.00 | 56.72 | 2.28 |
| 31589 | <13 y ND | 116 | 96 | 2 | 214 | 54.21 | 44.86 | 0.93 |
| 48588 | <13 y ND | 470 | 575 | 22 | 1067 | 44.05 | 53.89 | 2.06 |
| E386 | ≥ 13 y T1D | 1 | 38 | 136 | 175 | 0.57 | 21.71 | 77.71 |
| SC57 | ≥ 13 y T1D | 3 | 12 | 7 | 22 | 13.64 | 54.55 | 31.82 |
| SC76 | ≥ 13 y T1D | 98 | 84 | 146 | 328 | 29.88 | 25.61 | 44.51 |
| 6228 | ≥ 13 y T1D | 15 | 80 | 1242 | 1337 | 1.12 | 5.98 | 92.89 |
| 6563 | ≥ 13 y T1D | 48 | 143 | 251 | 442 | 10.86 | 32.35 | 56.79 |
| 6551 | ≥ 13 y T1D | 13 | 85 | 166 | 264 | 4.92 | 32.20 | 62.88 |
| 6520 | ≥ 13 y T1D | 108 | 181 | 472 | 761 | 14.19 | 23.78 | 62.02 |
| 6362 | ≥ 13 y T1D | 2 | 147 | 920 | 1069 | 0.19 | 13.75 | 86.06 |

| | | | | | | | | |
|---|---|---|---|---|---|---|---|---|
| 6550 | ≥ 13 y T1D | 7 | 91 | 179 | 277 | 2.53 | 32.85 | 64.62 |
| 6579 | ≥ 13 y T1D | 2 | 24 | 510 | 536 | 0.37 | 4.48 | 95.15 |
| 6469 | ≥ 13 y T1D | 5 | 71 | 1358 | 1434 | 0.35 | 4.95 | 94.70 |
| 6501 | ≥ 13 y ND | 664 | 480 | 149 | 1293 | 51.35 | 37.12 | 11.52 |
| 6374 | ≥ 13 y ND | 748 | 488 | 106 | 1342 | 55.74 | 36.36 | 7.90 |
| 6548 | ≥ 13 y ND | 232 | 89 | 19 | 340 | 68.24 | 26.18 | 5.59 |
| 6339 | ≥ 13 y ND | 488 | 642 | 157 | 1287 | 37.92 | 49.88 | 12.20 |
| 6431 | ≥ 13 y ND | 461 | 444 | 129 | 1034 | 44.58 | 42.94 | 12.48 |
| 6271 | ≥ 13 y ND | 1014 | 981 | 468 | 2463 | 41.17 | 39.83 | 19.00 |
| 6232 | ≥ 13 y ND | 366 | 664 | 216 | 1246 | 29.37 | 53.29 | 17.34 |
| 6333 | ≥ 13 y ND | 264 | 305 | 38 | 607 | 43.49 | 50.25 | 6.26 |
| 33366 | ≥ 13 y ND | 108 | 129 | 14 | 251 | 43.03 | 51.39 | 5.58 |
| 14666 | ≥ 13 y ND | 81 | 136 | 1 | 218 | 37.16 | 62.39 | 0.46 |
| PAN8 | ≥ 13 y ND | 321 | 185 | 41 | 547 | 58.68 | 33.82 | 7.50 |

Abbreviations: < 13 y ND, < 13 years without diabetes; < 13 y T1D, < 13 years at type 1 diabetes diagnosis; ≥ 13 y ND, ≥ 13 years without diabetes; ≥ 13 y T1D, ≥ 13 years at type 1 diabetes diagnosis.

### ESM Figures

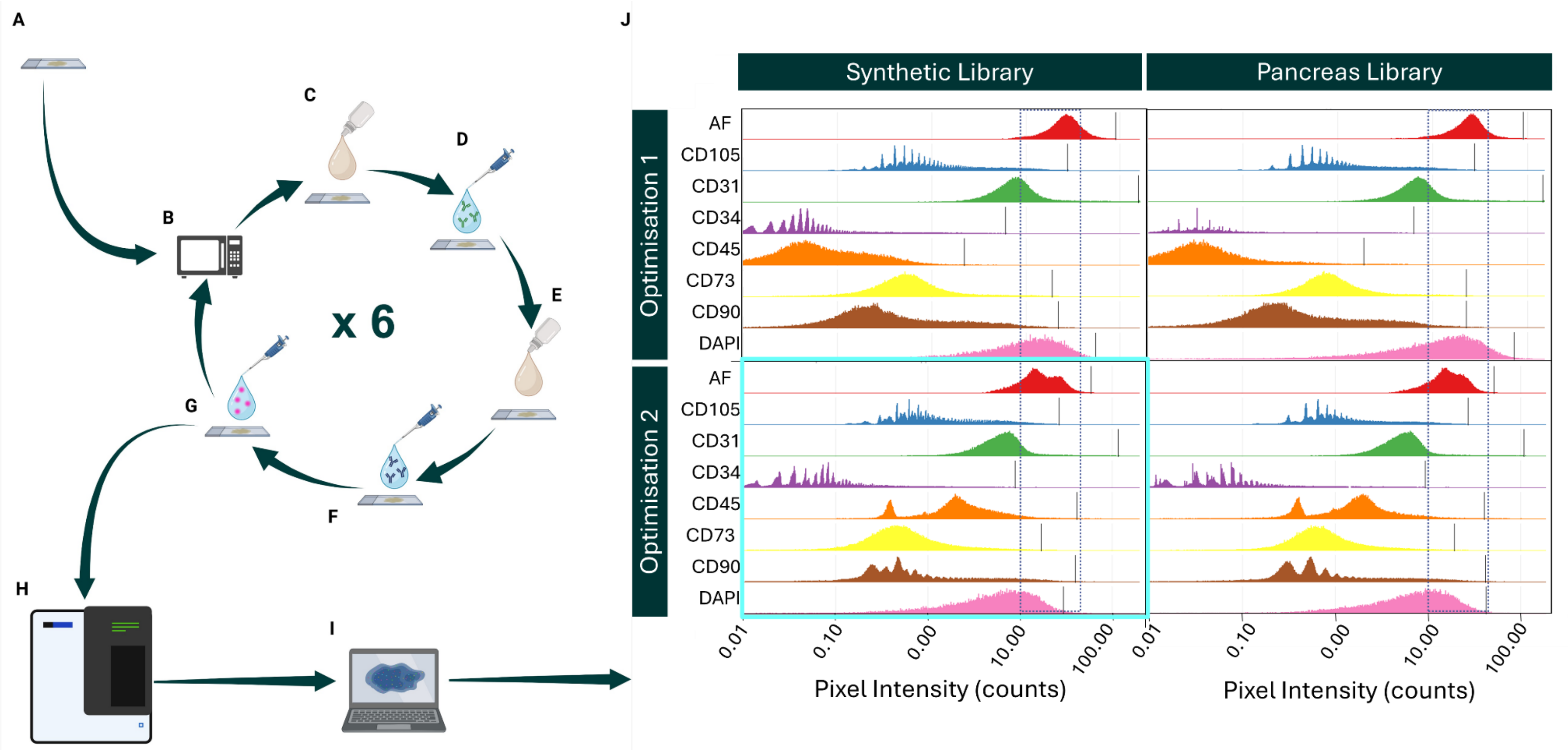


**ESM Figure 1** Optimisation of Opal-Tyramide signal amplification 6-plex immunohistochemistry to identify npMSCs.

Tissue sections (A) were subjected to 6 rounds of opal staining including heat-induced epitope retrieval (B), blocking (C), primary antibody incubation (D), endogenous peroxidase blocking (E), host-specific horseradish peroxidase-conjugated secondary antibody incubation (F), and signal generation (G). Whole-slide scans were acquired using the PhenoImager HT (Akoya Biosciences; H).

Six-plex opal immunohistochemistry required optimisation (J) before a final protocol could be confirmed. Optimisation was first performed (optimisation 1) with staining conditions informed by Akoya database of antigen clone-specific positioning experiments and our previous

experiments, in EADB pancreas and positive control tissue. Single-stained library and autofluorescence slides were also prepared with EADB pancreas tissue.
Images were unmixed using a custom pancreas library and the Akoya-provided synthetic library for comparison. PhenoptrReports Component Levels Report was performed in RStudio to investigate staining intensity (pixel intensity) and signal balance between fluorophores. A second optimsation was performed to refine and balance fluorophore signal intensity. Vertical lines along the histogram for each antigen are representative of the 99.9% percentile pixel and are representative of positive signal. The dashed rectangle shows optimal pixel intensity.
There was little difference between the custom pancreas library and the Akoya synthetic library in optimisation 2. Therefore, in the interest of pancreas tissue preservation, the synthetic library was chosen. Light blue box highlighting Optimisation 2, Synthetic library data shows the methodology confirmed as the final protocol.
Donor ID for tissue required for 6-plex opal and library/autofluorescence optimisation: PAN1, P39/67, E560, 8503, 6771/86, 88/66. Data shown in this figure from 88/66.
Figure creating using BioRender.

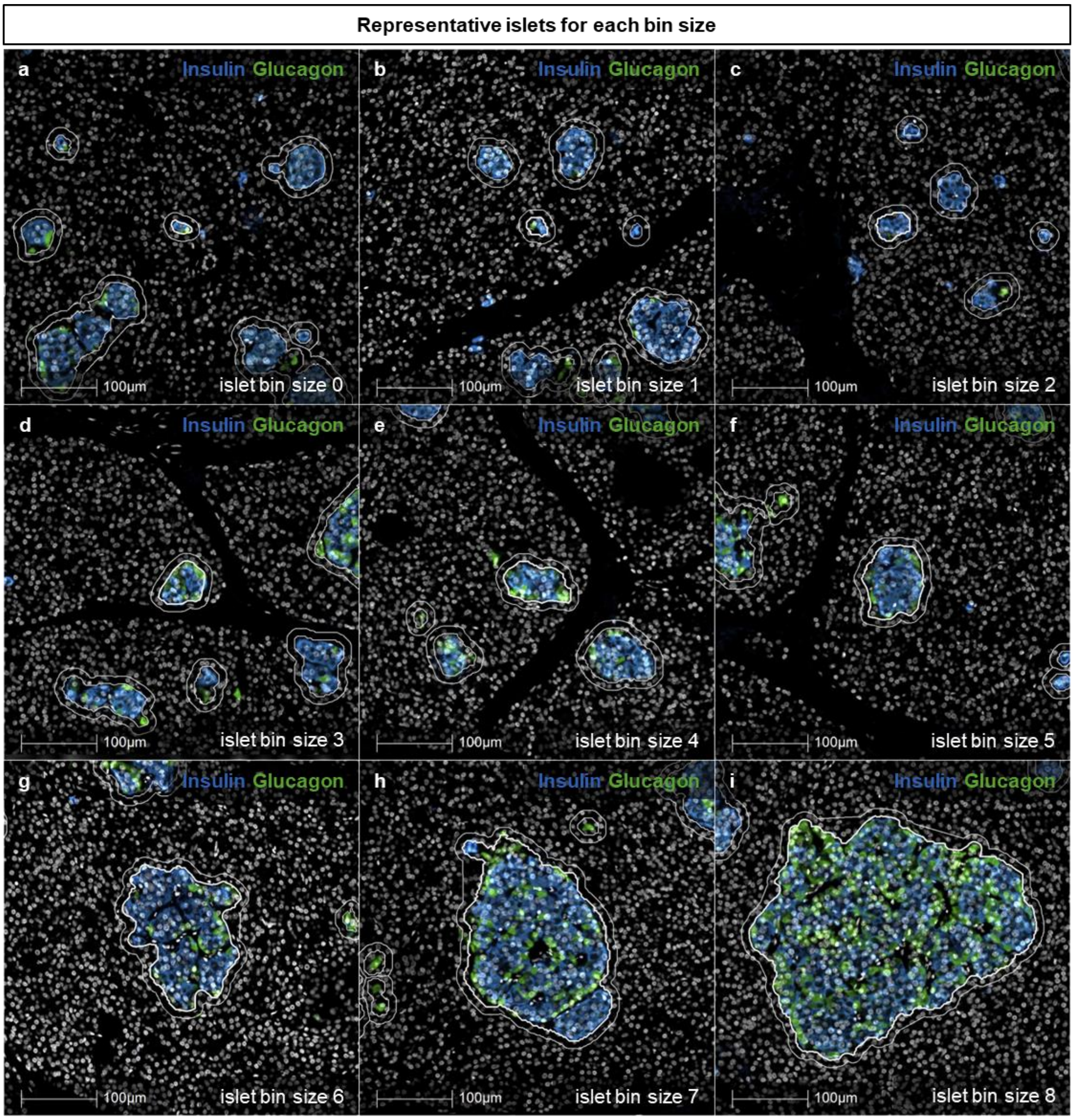


**ESM Figure 2** Representative islets from each islet bin size category.
White annotations show islets and islet periphery (10 µm outside of the islet), the bold white annotation in each micrograph highlights the islet which corresponds to the listed islet bin size. Donor ID: 6501, Individual ≥ 13 years without type 1 diabetes, nPOD.

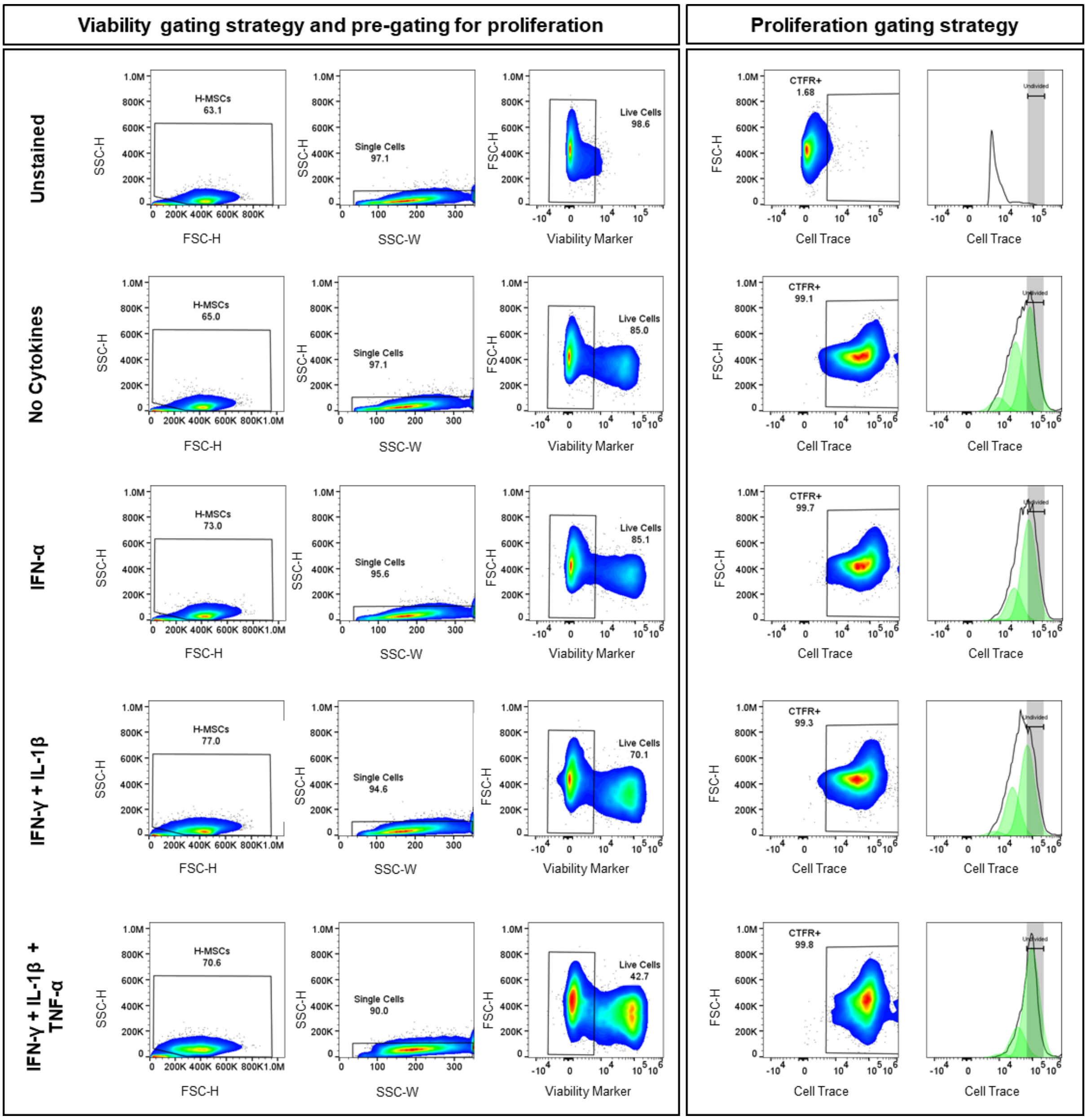


**ESM Figure 3** Gating strategy for in vitro viability and proliferation of cytokine-exposed MSCs. All representative figures from one independent experiment following three days cytokine exposure. Grey highlighted area within proliferation gating section represents gating for the undivided cell population.

Abbreviations: CTFR, CellTrace far red; FSC-H, forward scatter height; H-MSC, human mesenchymal stromal cell; SSC-H, side scatter hight; SSC-W, side scatter width.

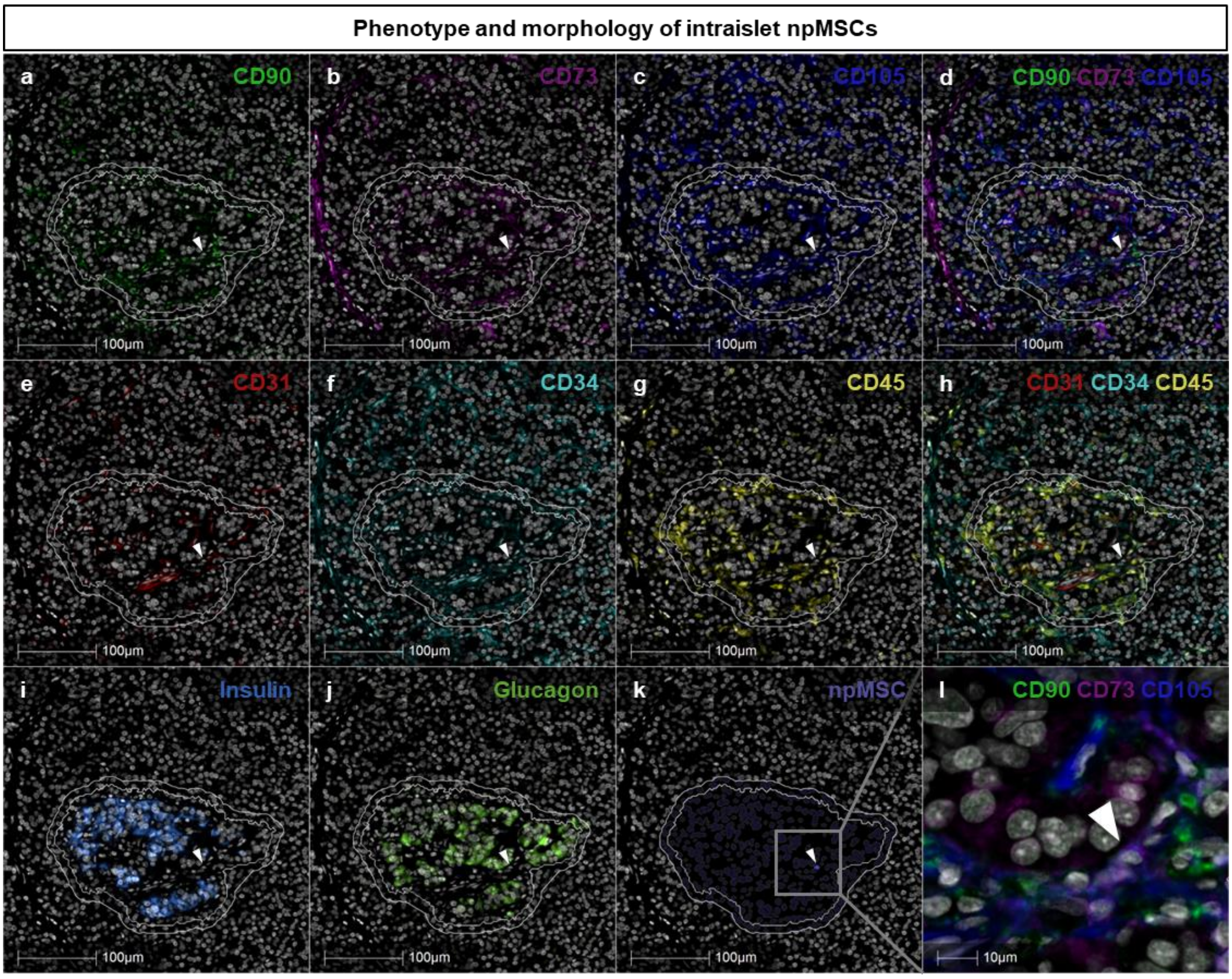


**ESM Figure 4** Phenotype and morphology of intraislet npMSCs.

Intraislet npMSCs (shown by white arrows; a-l) were identified in HALO as CD90+ (a), CD73+ (b), CD105+ ((c); overlay of positive markers, (d)), CD31- (e), CD34- (f), CD45- ((g); overlay of negative markers, (h)). Islets were identified by insulin (i) and glucagon (j) immunostaining (inner white annotation) and the islet annotation expanded 10 µm or until another annotation was reached (outer white annotation). Next, npMSCs were quantified (k). A magnified micrograph of k is shown in l (grey box shows magnified area) and intraislet npMSCs were identified (l).

Micrographs used in the representative figure were adjusted to optimise contrast and visibility without altering the underlying image data or quantification. Adjustments were made to the 'Black In', 'White In' and 'Gamma' settings.

Donor ID: < 13 years at type 1 diabetes diagnosis, 6578, nPOD.

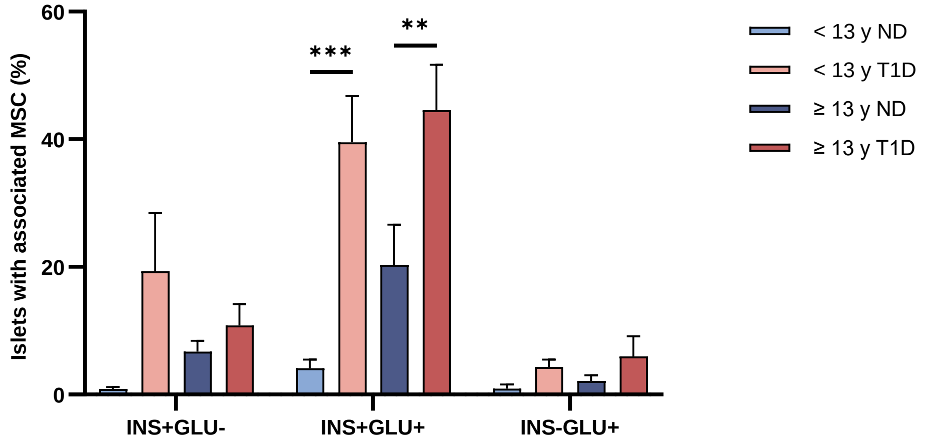


**ESM Figure 5** The proportion of islets with an associated npMSC is increased for insulin-containing glucagon-containing islets in type 1 diabetes.

Islets with one or more npMSC either inside the islet or within 10 μm of the islet periphery were defined as having an associated npMSC.

Data derived from *n* = 26,376 individual islets from 38 individuals (*n* = 8 individuals < 13 years at type 1 diabetes diagnosis and *n* = 11 individuals ≥ 13 years at type 1 diabetes diagnosis; *n* = 8 individuals < 13 years without diabetes and *n* = 11 individuals > 13 years without diabetes). Bars represent mean ± SEM. Two-way ordinary ANOVA with post hoc tests adjusted for multiple comparisons with Bonferroni correction. Post hoc comparisons: type 1 diabetes group and age-matched individuals without diabetes; type 1 diabetes and age of diagnosis; individuals without diabetes. Comparisons shown with a horizontal black line between the two compared groups.

***$p \leq 0.001$; **$p \leq 0.01$

Abbreviations: < 13 y ND, < 13 years without diabetes; < 13 y T1D, < 13 years at type 1 diabetes diagnosis; ≥ 13 y ND, ≥ 13 years without diabetes; ≥ 13 y T1D, ≥ 13 years at type 1 diabetes diagnosis.
